# Macroalgal fucoidan can activate the biological carbon pump

**DOI:** 10.64898/2026.07.10.737761

**Authors:** Inga Hellige, Hagen Buck-Wiese, Margot Bligh, Timothy Thomson, Lydia White, Carol Arnosti, Dariya Baiko, Linda Biehler, Mar Fernández-Méndez, Sherif Ghobrial, Bowei Gu, Camilla Gustafsson, Mohammed Kajee, C. Chad Lloyd, Nguyen P. Nguyen, Miriam Philippi, Dan Potin, Philippe Potin, Mark Rothman, Beatriz S. Murillo, Michael Seidel, Catharina Uth, Evie Wieters, Marie Magnusson, Jan-Hendrik Hehemann

**Author notes:** These authors contributed equally to this work (Inga Hellige, Hagen Buck-Wiese).

## Abstract

Macroalgae secrete complex carbohydrate polymers, their extracellular matrix, as protection against microbial degradation. By resisting breakdown, these carbohydrates can contribute to marine carbon sequestration, though mechanisms, extent, and timescales remain unknown. Using ship-based sampling and experiments, we found that brown macroalgae release 1.7-4.2% of carbon fixation as fucoidan, equivalent to 0.32-0.88 mg fucoidan per gram of dry seaweed tissue per day. A Bayesian model trained on our empirical data, coupled with Monte Carlo simulations suggests an annual global release of 13-37 megatons fucoidan carbon. Moreover, degradation resistance combined with surface-activity enabled fucoidan to act as glue that cross-linked allochthonous organic carbon including microbes and proteins into marine snow. Notably, substantial fucoidan exudation was universally conserved across all tested species and regions. Thus, any brown macroalgal species can be used e.g. via aquafarming to enhance the formation of marine snow.

## Main

Remnants of a molecular arms race in the sunlit ocean bury carbon for centuries in the ocean floor^1^. This arms race starts with diverse photosynthetic algae, which collectively fix about 45 gigatons of carbon per year^2^. Rapid carbon dioxide fixation enables algae to cover their cells with secreted carbohydrate polymers, or glycans^3^. Their diverse structures evolve to maintain a physicochemical barrier, the extracellular matrix, which protects against recognition and enzymatic invasion by pathogens including viruses and heterotrophic bacteria^4^. Thus algal glycans and bacterial proteins are the weapons of this molecular arms race. Bacteria can degrade even the most structurally diverse glycans with carbohydrate-active-enzymes to completion under optimal conditions in the laboratory^5,6^. However, ship-based studies reveal an abundance of glycans indicating that the ocean is a suboptimal environment for bacterial glycan degradation^7,8^. For example, radiocarbon dating shows ∼10 Gt carbon, of dissolved glycans, accumulate for up to a decade (average age ∼3 years) in surface seawater^9,10^. In laboratory experiments algal glycans also persist but at higher concentrations (>1 mg/L,^11,12^). Turbulent conditions increase the encounter rate of dissolved glycans with other, hitherto unidentified molecules, promoting co-precipitation and thereby lowering their concentration in the ocean^13–15^. This surface-active behavior acts as a nucleus for the formation of marine snow, which transports organic carbon, including degradation-resistant glycans, to the ocean floor^16–18^. About 90% of this carbon export is confined to coastal margins, where algal glycans are buried and persist in sediments for centuries^19–24^. However, which glycans drive this process - known as the biological carbon pump^25^ - remains unknown. This lack of knowledge stems from the fact that during the past decades mainly the sum of glycan carbon has been quantified in marine organic matter without the molecular resolution required to resolve their many different individual structures, quantities, and sources^26^. For example, ∼ 1 % of the marine dissolved organic matter pool (∼650 Gt C), consists of polymeric fucose of unknown origin^10^.

One of the most photosynthetically active marine phyla of algae^2^, the ochrophytes, including brown macroalgae and microalgae (e.g. kelp, diatoms), which are important for marine carbon sequestration^27–29^, secrete as part of their extracellular matrix fucoidan^30–33^. Fucoidan is an umbrella term for a structurally diverse family of extensively sulfated fucans. The main chain usually consists of alpha-configured, 1,3- or 1,4-linked fucose, which can be decorated with species specific side chains composed of other monosaccharides. In some species e.g. the pelagic macroalgae *Sargassum* spp. and the microalgae *Glossomastix* spp. the main chain contains other monomers besides fucose^12,34^. Additionally, fucoidan shows structural hallmarks of an ongoing adaptive radiation, which manifests in a broad-acting, anti-cellular function^35^. This diversification, extensive sulfation, and anti-cellular function render fucoidan particularly resistant against microbial degradation^36^.

We previously hypothesized that brown macroalgae exude fucoidan^37^ and that this macromolecule could potentially be used for negative carbon dioxide emissions^38^ by acting as glue that cross-links and precipitates other organic molecules in the ocean^26^. Here, we report on the previously undocumented scale and geographical distribution of fucoidan exudation across the phylum of marine macroalgae. Moreover, we uncovered the interaction with and precipitation of proteins as a previously unknown molecular mechanism required for marine snow formation.

### Fucoidan presence in coastal surface waters

First, we tested whether natural and farmed populations of brown macroalgae release dissolved fucoidan into the ocean (**Fig. 1, Table S1, Fig. S1**). Natural populations were investigated in South Africa, New Zealand, France, Chile, Finland, the western North Atlantic, and Mexico. The studied *Saccharina latissima* aquafarm was located in Northern Ireland. Fucoidan was quantified using previously described methods^37^ with detailed descriptions in SI methods. Briefly, to quantify total dissolved fucoidan we desalted seawater samples using dialysis, retaining molecules >1 kDa. Samples were hydrolyzed with acid to release monosaccharides, which were quantified via chromatography with pulsed amperometric detection. Fucoidan is the only known marine glycan family primarily composed of fucose^39^. Other sources of fucose, aside from mammalian milk oligosaccharides, are glycoproteins and bacterial exopolysaccharides but they contain only a fraction of fucose^40,41^. Therefore, and because fucose is the only monomer that is shared across diverse fucoidan types, we used polymeric fucose as a quantitative proxy for dissolved fucoidan^37,42^. A second method targeted dissolved surface active fucoidan (SAF), using anion exchange chromatography (AEX) to extract only negatively charged fucoidan from seawater prior to acid hydrolysis and quantification.

**Figure 1.**
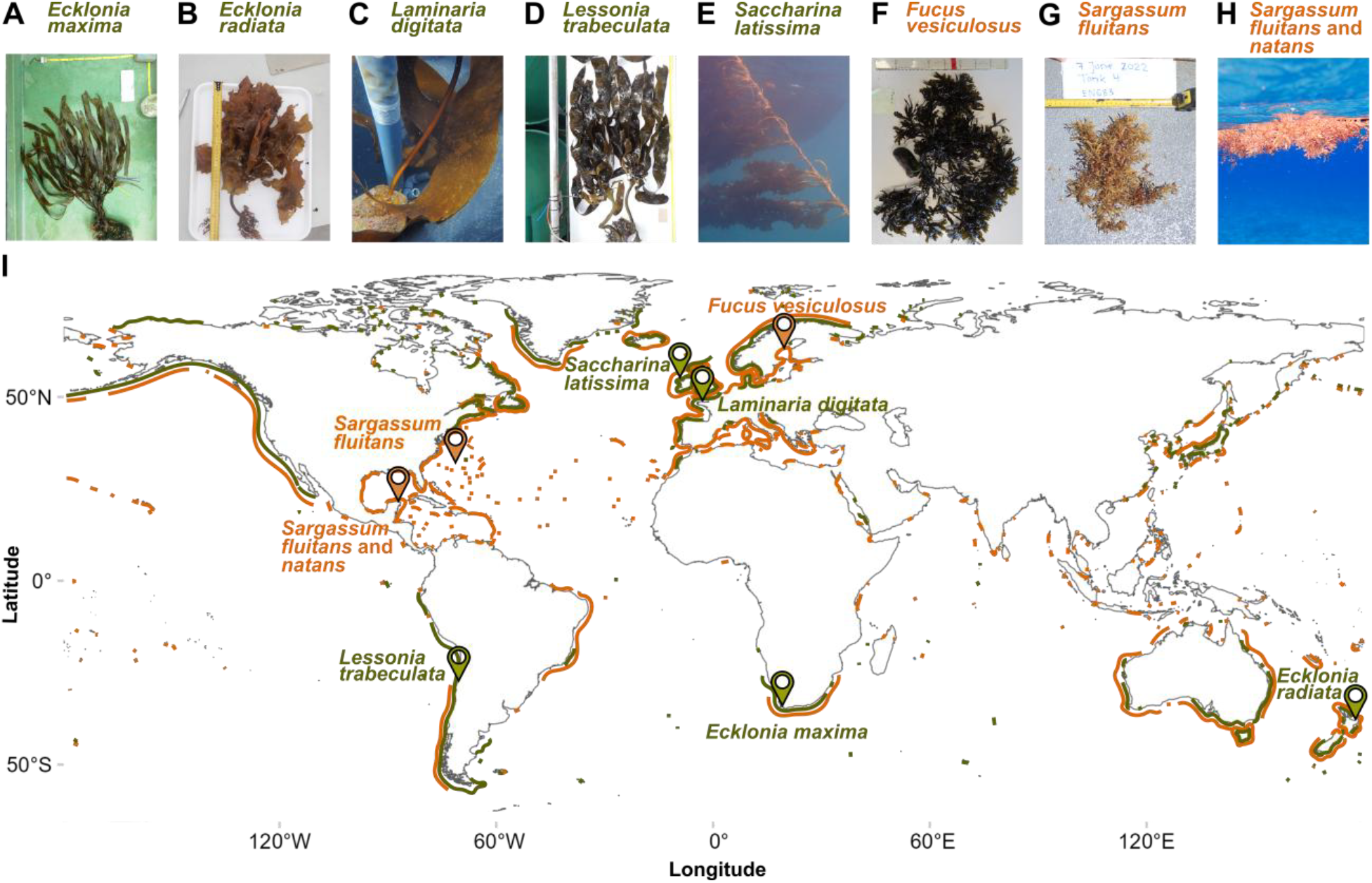
Sampling representatives of a global taxon and their surroundings. **A-H)** Images of sampled algae and **I)** occurrence of brown algae based on a recent distribution dataset^43^ with markers indicating sampling locations. Distributional lines correspond to Laminariales (green) and Fucales (orange).

We detected 0.02-0.09 mg L^-1^ SAF, with higher in-situ values in close spatial proximity to algal populations, suggesting these species release fucoidan into the ocean (**Fig. S1**). Fucoidan concentrations increased towards macroalgae for transects in South Africa, France, and Mexico (**Fig. S1A, C, G**). The highest SAF concentrations of 0.093 ± 0.003 mg L^-1^ (mean ± s.e.m) and 0.075 ± 0.015 mg L^-1^ were detected inside a patch of floating *Sargassum spp*. off the coast of southern Mexico (**Fig. S1G**) and in outgoing tidal surface waters downstream of *L. digitata* beds in France (**Fig. S1C**), respectively. Generally, SAF concentrations were elevated in surface waters inhabited by brown macroalgae and rapidly decreased with depth. This distribution supports the hypothesis that brown macroalgae are the main source of fucoidan in their respective water layers (**Fig. S1**).

### Release and persistence of dissolved fucoidan

To quantify net release rates, we conducted mesocosm experiments with brown macroalgal species dominant to, and locally sourced from, their respective environments (**Fig. 1, S2, Table S2**). Triplicate mesocosms received one brown macroalgal specimen and local seawater, including microbial communities. Triplicate control mesocosms without macroalgae received the “same” seawater. Mesocosms were exposed to a natural photoperiod cycle and supplied with air stones. Fucoidan concentrations were measured at two-day intervals for the duration of the 24-day experiments, with macroalgae removed on day 12 (**Fig. 2**). Until day 12, mesocosm water was periodically replenished with fresh seawater to replace potentially depleted nutrients (**Fig. S3A**). Throughout the incubation period, macroalgae maintained primary productivity, which scaled with biomass (**Fig. 2A**). Accounting for water replenishment, fucoidan concentrations peaked on day 12, when macroalgae mesocosms reached 1.52 ± 0.32 mg L^-1^ dissolved fucoidan and 0.64 ± 0.11 mg L^-1^ SAF compared to mesocosms without macroalgae at 0.29 ± 0.07 mg L^-1^ dissolved fucoidan and 0.21 ± 0.08 mg L^-1^ SAF (**Fig. 2B, C**).

**Figure 2.**
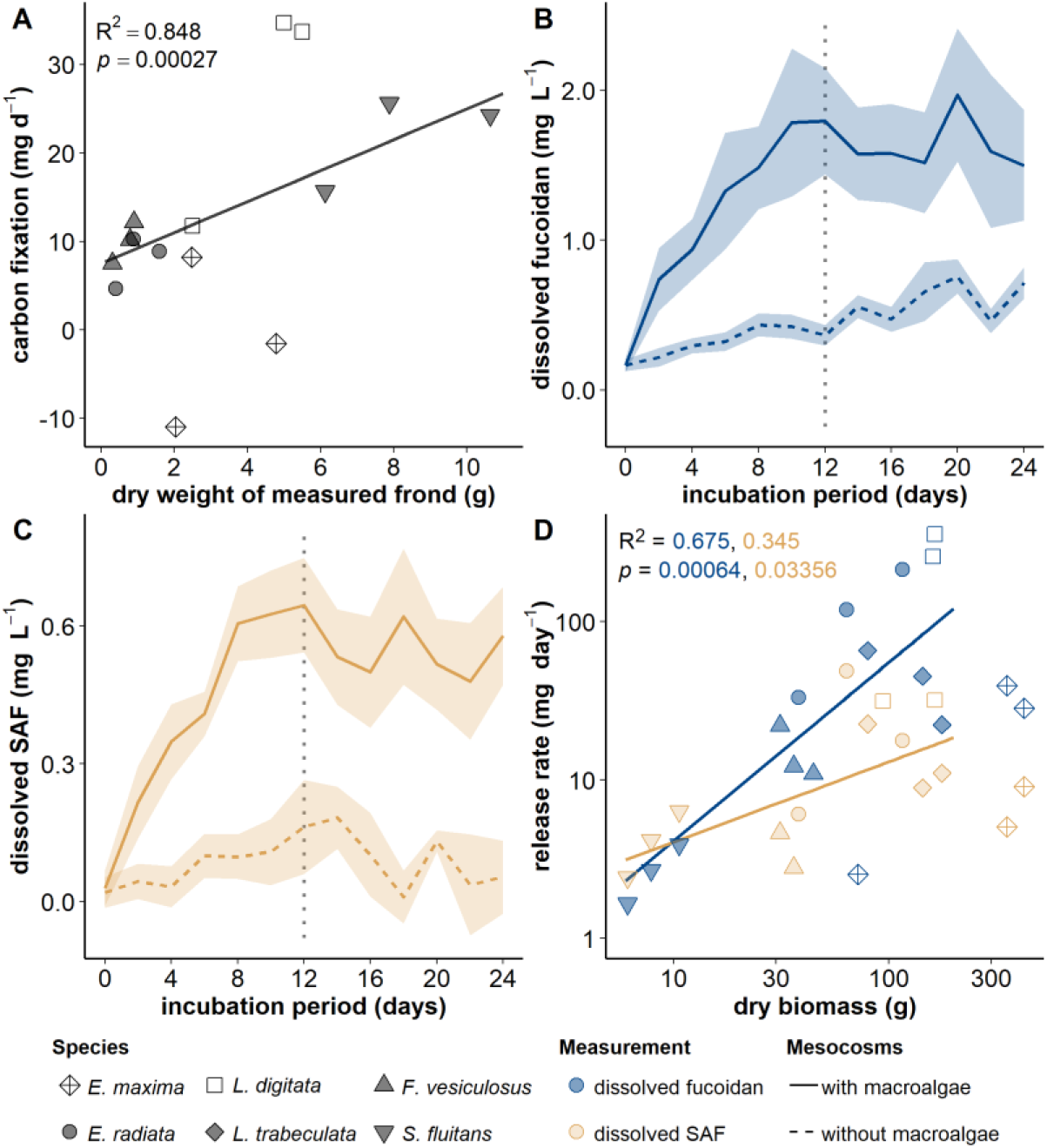
Release of fucoidan by brown macroalgae scales with biomass. **A)** Net primary production plotted against biomass of measured frond, with regression based on log10-transformed values and excluding *E. maxima* due to insufficient light and *L. digitata* due to missing water exchange (n = 9). No data available for *L. trabeculata*. **B-C)** Cumulative fucoidan concentrations in mg L^-1^ of macroalgae mesocosms (solid line) and mesocosm without macroalgae (dashed line) over 24 days of incubation, excluding *L. digitata* and *E. maxima* due to experimental limitations, removal of macro algae shown by vertical dashed grey line **B)** total dissolved fucoidan, **C)** dissolved surface active fucoidan (SAF). **D)** Release of total fucoidan (blue) and SAF (yellow) during the first four days of experiment plotted against dry biomass of macroalgae. Regression models in D use log10-transformed values and exclude *E. maxima* and *L. digitata* (n = 12).

Fucoidan detected in mesocosms without macroalgae is consistent with diatom microalgae producing a fucoidan variant also referred to as fucose-containing sulfated polysaccharide (FCSP)^26^. In the *E. radiata* mesocosms, removing microalgae by adding filtered seawater from day 2 onwards, caused the fucoidan synthesis signal to disappear in treatments without macroalgae. (**Fig. S4H, N**). Based on a hierarchical Bayesian model, macroalgae mesocosms had 0.52 mg L^-1^ higher dissolved fucoidan (95% confidence interval: 0.31-0.72 mg L^-1^) and 0.18 mg L^-1^ higher SAF (0.13-0.22 mg L^-1^) concentrations than mesocosms without macroalgae (see supplementary section **Bayesian models**).

Next, we calculated the net release rate as a function of algal biomass and in relation to net primary productivity. On average, released fucoidan carbon accounted for 1.7-4.2% of net primary productivity, which correlated with dry weight whilst displaying inter-specific variability (**Table 1, Fig. 2D, S5, S6**; for details see supplementary section **Primary productivity**). The release of both dissolved fucoidan and SAF correlated significantly with macroalgal biomass (*p* < 0.001, R^2^ = 0.67 for dissolved fucoidan and *p* = 0.034, R^2^ = 0.35 for SAF) (**Fig. 2D**). These results demonstrate that the quantity of brown macroalgal biomass present in the environment is a major factor underpinning the rate at which fucoidan is synthesized and released into the water column.

**Table 1.** **Fucoidan release rates** based on accumulation during the first four days of incubation in the dissolved fucoidan and the dissolved surface active fucoidan (SAF) fraction. Reported values include release per volume and time (mg L^-1^ d^-1^), per unit brown algae biomass and time normalized by mesocosm volumes (mg g_dw_^-1^d^-1^) and percentage of net primary production (NPP) detected as released fucoidan carbon.

|  | brown algae | *E. maxima* | *E. radiata* | *L. digitata* | *L. trabeculata* | *F. vesiculosus* | *S. fluitans* | mean |
| --- | --- | --- | --- | --- | --- | --- | --- | --- |
| incubation | dry weight (g) | 280.8 ± 68.0 | 79.5 ± 15.5 | 139.5 ± 14.4 | 133.4 ± 17.9 | 37.4 ± 2.5 | 8.2 ± 0.8 |  |
|  | volume (L) | 220 | 220 | 250 | 250 | 57 | 40 |  |
| dissolved fucoidan | accumulation (mg L<sup>-1</sup> d<sup>-1</sup>) | 0.17 ± 0.04 | 0.58 ± 0.24 | 1.43 ± 0.34 | 0.33 ± 0.15 | 0.36 ± 0.06 | 0.09 ± 0.01 | 0.49 ± 0.20 |
|  | accumulation (mg g<sub>dw</sub><sup>-1</sup> d<sup>-1</sup>) | 0.13 ± 0.01 | 1.63 ± 0.29 | 2.27 ± 0.4 | 0.89 ± 0.50 | 0.58 ± 0.12 | 0.42 ± 0.04 | 0.98 ± 0.33 |
|  | % of NPP | negative NPP | 9.85 ± 3.69 | 5.94 ± 1.22 | no NPP measured | 0.57 ± 0.13 | 2.46 ± 0.17 | 4.23 ± 1.05 |
| dissolved SAF | accumulation (mg L<sup>-1</sup> d<sup>-1</sup>) | 0.04 ± 0.02 | 0.12 ± 0.06 | 0.18 ± 0.07 | 0.04 ± 0.02 | 0.06 ± 0.02 | 0.17 ± 0.03 | 0.10 ± 0.03 |
|  | accumulation (mg g<sub>dw</sub><sup>-1</sup> d<sup>-1</sup>) | 0.02 ± 0.01 | 0.32 ± 0.19 | 0.36 ± 0.19 | 0.11 ± 0.06 | 0.11 ± 0.04 | 0.75 ± 0.05 | 0.28 ± 0.11 |
|  | % of NPP | negative NPP | 2.41 ± 1.81 | 0.95 ± 0.46 | no NPP measured | 0.11 ± 0.04 | 4.45 ± 0.52 | 1.78 ± 0.52 |

Mesocosm experiments on locally dominant species consistently demonstrated the release of fucoidan by brown macroalgae (**Table S2**) in six contrasting marine regions, including eutrophic, coastal upwelling, and open ocean environments (**Fig. 1, S1**). We measured dissolved organic carbon (DOC) concurrently and found dissolved fucoidan accounted for 30-50% and SAF for 15-25% of DOC accumulation, respectively (**Fig. S4, S7**). This accumulation is consistent with structurally diverse dissolved glycans, including diatom fucoidan, are stable for month-to-years in the ocean^9,26^. We noted diminishing fucoidan accumulation rates over the course of the experiments (**Fig. S8**). This decrease suggested precipitation may have removed the fucoidan from solution^15,17^. Hence, we tested whether macroalgal fucoidan is also stable enough to persist and aggregate in the presence of heterotrophic microbes into degradation resistant particles.

To assess the potential stability of fucoidan under laboratory conditions, we monitored concentrations within water collected from mesocosm experiments on day 24, held in 5-10 L containers for 12 months. Incubations were performed in the dark to avoid new photosynthetic production. We detected dissolved fucoidan throughout the sampling period but observed a decrease in concentrations over time (**Fig. S9, S10, S11**). Samples collected from the macroalgae mesocosms had 0.29 mg L^-1^ (-0.07-0.66 mg L^-1^) higher dissolved fucoidan concentrations compared to samples from control mesocosms without macroalgae after 9 and 12 months, based on a hierarchical Bayesian model with treatment as predictor and location as random factor (**Fig. S11**). Fucoidan-specific monoclonal antibodies, which showed successful binding to fucoidan extracted from *L. trabeculata* biomass, confirmed the persistence of dissolved fucoidan in *L. trabeculata* mesocosms (**Fig. S12D**). In contrast, SAF had decreased to values close to the detection limit (**Fig. S11**), suggesting selective removal of this fraction from the dissolved pool. Preferential degradation of the SAF fraction by bacteria was unlikely, as sulfation generally enhances recalcitrance^36,44,45^. Yet, anionic glycans are highly surface-active, which promotes precipitation from dissolved into particulate organic matter^15,26,46,47^.

### Brown macroalgal fucoidan precipitates organic matter

To test precipitation as a potential removal process, we quantified particulate organic matter fractions during our 24 day-long mesocosm experiments. Particulate organic carbon increased significantly over time in mesocosms with macroalgae, including after the algae were removed (linear model on log_10_-transformed data, *p* = 0.028, R^2^ = 0.07), in contrast to mesocosms without macroalgae (*p* = 0.934; **Fig. S13A**). Similarly, the amount of particulate fucoidan increased only in macroalgae mesocosms (linear model on log_10_-transformed data, *p* < 0.001, R^2^ = 0.27), while remaining unchanged in mesocosms without (*p* = 0.959; **Fig. 3A, S13B, S14**). Notably, particulate fucoidan also increased after the removal of macroalgae, hinting toward the precipitation of dissolved fucoidan, rather than shedding of algal detritus. In the nutrient-rich upwelling regions with *E. maxima* (**Fig. S14A**) and *L. trabeculata* (**Fig. S14D**), fucoidan-containing particles were abundant in all mesocosms. A similar pattern was observed for *L. digitata* mesocosms, where the added seawater contained background particulate fucoidan concentrations of up to 0.17 mg L^-1^, which were responsible for the lack of contrast between background and newly formed particles (**Fig. S14C**). The highest contrast arose in oligotrophic Sargasso Sea incubations, where fucoidan-containing particles were exclusively present in mesocosms with *S. fluitans* (**Fig. S14F**).

**Figure 3.**
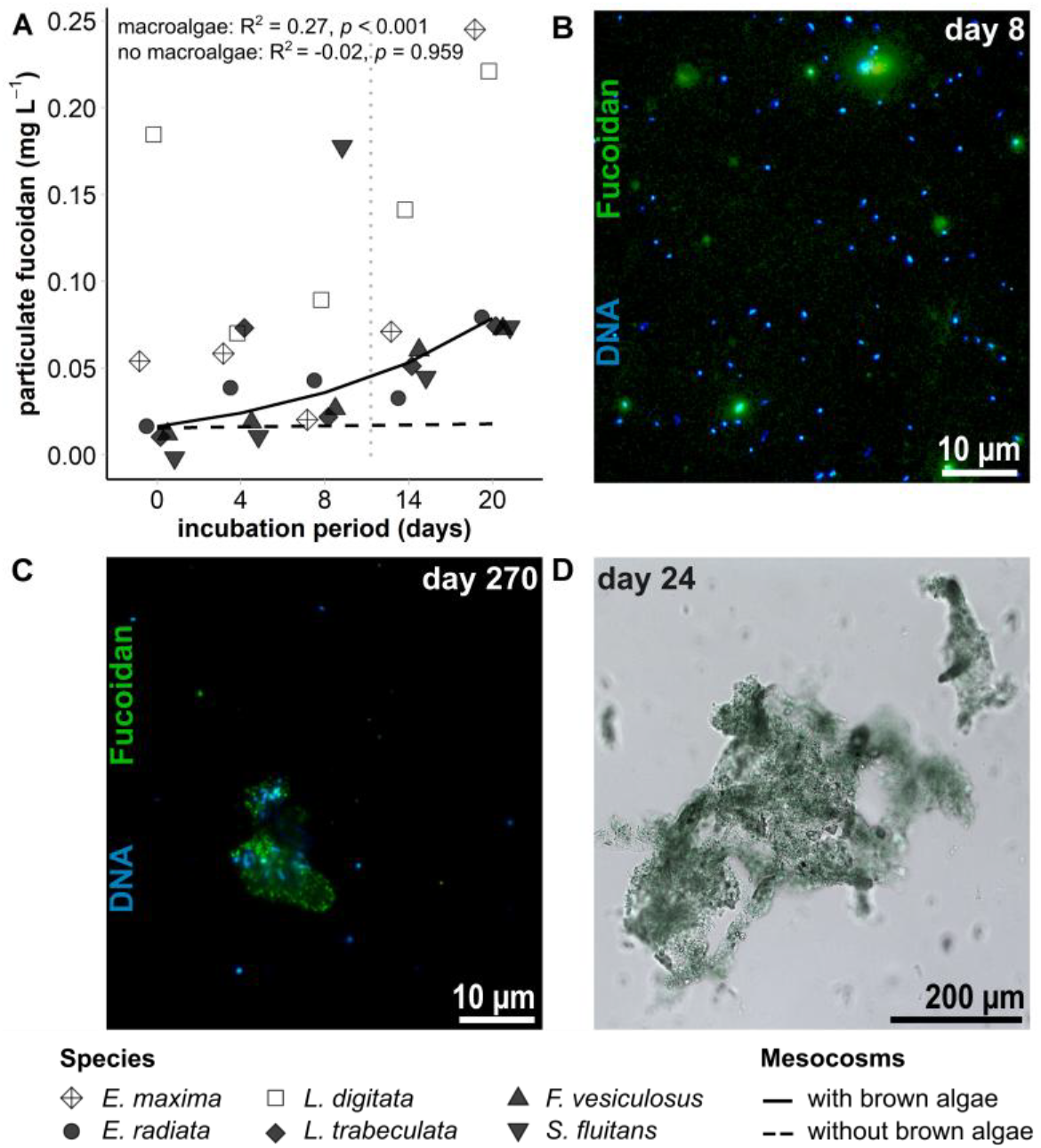
Brown algal fucoidan precipitates into particles that can persist for months. **A)** Total particulate fucoidan in mg L^-1^ estimated from acid-released fucose in mesocosms with macroalgae (black, solid line) and without macroalgae (dashed line), with lines indicating linear models of log^10^-transformed concentrations, removal of macroalgae shown by vertical dashed grey line. Calculations exclude *E. maxima* and *L. digitata*. Individual data points are only shown for mesocosms with macroalgae to enhance clarity, all data is displayed in **Fig. S13B. B-C)** Filters from *F. vesiculosus* mesocosms showing DAPI-stained cells (blue) and BAM1 immunolabeled fucoidan (green) for **B)** day 8 and **C)** day 270 in the dark. **D)** Brightfield microscopic image of precipitate after 24 days at bottom of *L. trabeculata* incubation.

Microscopy of filters probed with the fucoidan-specific BAM1 antibody confirmed the precipitation of fucoidan into recalcitrant particles and revealed precipitation with exogenous matter, including DAPI-stained microbial cells. (**Fig. 3B-C, S15**). *F. vesiculosus* incubations, which we sampled for particle imaging, were enriched in BAM1-stained particles after 8 days (**Fig. 3B**) and screening filters for particles after 9 months of incubation in the dark showed that fucoidan precipitation had continued, resulting in formation of persistent and larger fucoidan-rich particles (**Fig. 3C**).

Microbial cells bound to precipitates with a matrix of fucoidan indicate precipitation with bacteria, which may have contributed to declines in bacterial cell abundances in the water column (**Fig. S14G-L**). Particulate fucoidan concentrations not only correlated with POC (**Fig. S16**), but in high-nutrient environments also with particulate nitrogen (e.g. *F. vesiculosus* in the Baltic: linear model, *p* < 0.001, R^2^ = 0.70) (**Fig. S17**). Sedimented particles from mesocosm incubations showed fucoidan antibody signal among a multitude of visible components (**Fig. 3D, S19**). Fucose comprised 3-12% of monosaccharides in the sedimented particles, in line with the precipitation of allochthonous carbon including microbial cells (**Fig. S19A-B**). The formation and month-long persistence of particles suggests that fucoidan promotes the stabilization and export of allochthonous carbon.

To evaluate whether this process also occurs under natural conditions, we examined fucoidan dynamics along a transect through a kelp farm. Samples from a transect through a kelp farm supported fucoidan precipitation in the environment, although fucoidan precipitation could not be distinguished from potential shedding of kelp detritus in our analysis (**Fig. S20**). SAF concentrations appeared elevated within the kelp farm (2.32 ± 0.79 mg L^-1^) and downstream (2.11 ± 0.41 mg L^-1^) compared to upstream (1.62 ± 0.48 mg L^-1^). Concentrations of particulate fucoidan increased by 1.06 mg L^-1^ (Bayesian model, 95% confidence interval: 0.06-2.05 mg L^-1^) downstream of the *S. latissima* farm compared to samples taken upstream of and inside the farm (**Fig. S20**).

Marine particles can form when polysaccharides spontaneously assemble into transparent exopolymer particles (TEP)^17,46^, the precursors of marine snow **(Fig. 4A)**. Although fucoidan is characterized by its high solubility of up to 10 g/L, concentrations far higher than those reached in our mesocosms, let alone in the environment, we nevertheless observed fucoidan forming precipitates with unknown nitrogen containing compounds (**Fig. S17E**). Thus, we considered proteins could be a partner molecule required for precipitation, as both are part of marine particles^48,49^. By estimating fucoidan from fucose concentrations in literature values and mesocosms without macroalgae, and assuming an average C content of 0.27 for fucoidan^37^, we found that the contribution of fucoidan C-to-POC ratio had a narrow distribution around 1.5 ± 0.1% across oceanic regions (linear models on log_10_-transformed data, *p* < 0.001 for both, R^2^ = 0.90 and 0.70, respectively). When brown macroalgae were present, however, the contribution reached up to 25%, albeit without significant correlation (**Fig. S18**). Thus, we combined fucoidan and protein in a 1:100 (w/w) ratio, which led to precipitation during 16 days of incubation in a roller tank, increasing molecular encounter rates. Fucoidan concentrations as low as 0.01 mg L^-1^ in the presence of 1 mg L^-1^ protein led to the emergence of 5,000-6,000 particles mL^-1^ within two days (**Fig. 4B**, yellow lines, grey symbols). In samples with 0.05 and 0.1 mg L^-1^ fucoidan, 50,000 particles mL^-1^ formed in 16 days. Importantly, without the respective binding partner the molecules remained soluble, showing that fucoidan and proteins are interacting agents of precipitation (**Fig. 4B**, yellow lines, yellow circles and grey lines, grey circle control treatments). Consistent with particle composition (**Fig. S18**), a ∼1% fucoidan fraction precipitated other molecules even at dilute concentrations. The precipitation of proteins is consistent with the spectral properties of “freshly” formed marine snow that appears, akin to coagulated proteins, “egg-white” **(Fig. 4A)**. Moreover, our results suggest that the stickiness of anionic glycans is context dependent for it requires the presence of other molecules. This feature is broadly known from and applied in common, household multi-component glues or in laboratory polyacrylamide gels that solidify only after mixing the different molecules. The tested concentrations and binding partners evidently only represent a subset of environmentally relevant conditions important to understand marine snow formation.

**Figure 4.**
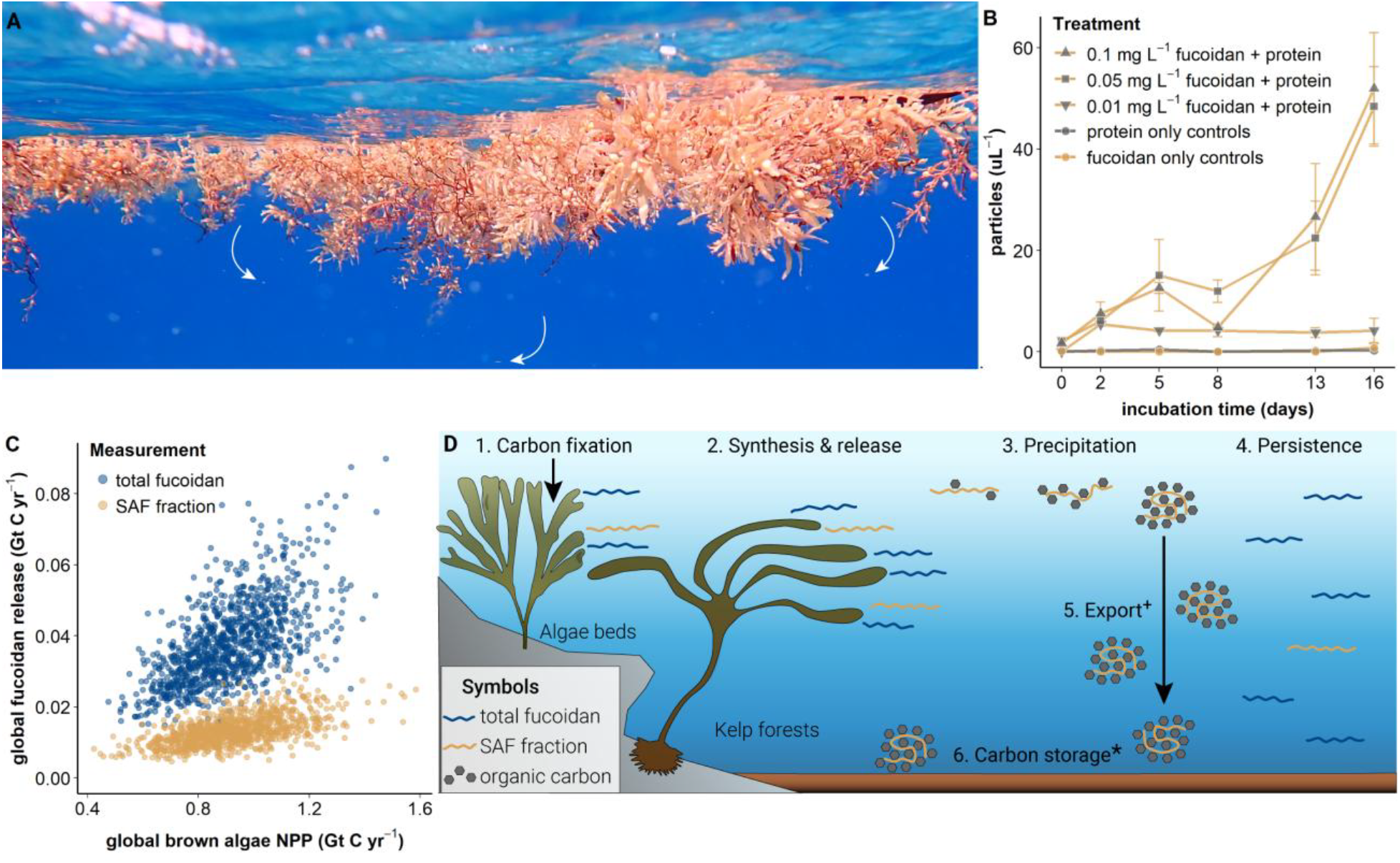
Fucoidan released by macroalgae drives organic matter precipitation. **A)** Holopelagic *Sargassum* spp. with marine snow particles highlighted by white arrows off the coast of Quintana Roo, Mexico, in July 2022. **B)** Particles counted via flow cytometry in roller tank incubations containing SAF fucoidan from *L. hyperborea* and the protein lysozyme (n = 3). In incubations with fucoidan, protein was supplied at a 1:100 (w/w) ratio, e.g. 0.01 mg L^-1^ fucoidan with 1 mg L^-1^ protein (yellow lines, grey symbols). Control incubations included the same protein concentrations without fucoidan (grey line, grey circles) and the same fucoidan concentrations without protein (yellow line, yellow circles). **C)** Modeled annual release of fucoidan carbon as dissolved fucoidan (blue) and SAF (yellow) fractions from mesocosm-based Bayesian models and net primary productivity estimates^50^. **D)** Conceptual illustration of the release of dissolved fucoidan and its environmental fate. **[1-4]** Surface processes documented in this work are stated at the top. **[5]** Export^**+**^ flux processes remain largely unconstrained. **[6]** Carbon storage* in the form of fucoidan has been shown on climate-relevant timescales (>100 years) in marine sediments^19–21^.

To estimate the amount of organic carbon that can precipitate via this process, we scaled fucoidan release rates based on macroalgal productivity^50^. First, we used a hierarchical Bayesian model approach with our mesocosm data to compute the amount of fucoidan released per biomass (supplementary section **Bayesian models**). Trained exclusively on the concentration change over time, our model isolated the macroalgae effect as elevating dissolved fucoidan by 0.26 mg g^-1^_dw_ d^-1^ and SAF by 0.09 mg g^-1^_dw_ d^-1^. Simulations based on the resulting posterior distributions of brown algae biomass and first order removal effects proved these two parameters sufficient to reproduce the experimentally observed trends (**Fig. S21**). The thereby validated posterior distributions served in conjunction with primary production rates (**Fig. 2A**) to extrapolate fucoidan release to global subtidal brown algal productivity treating biomass as the shared driver in Monte Carlo simulations^50^. Based on a global carbon fixation rate of 917 Mt yr^-1^, the simulations projected an annual release of total fucoidan carbon (**Fig. 4C**, blue circles) with a mean of 37 Mt yr^-1^, and thereof SAF (**Fig. 4C**, yellow circles) with a mean of 13 Mt yr^-1 50^. Considering the ratio at which SAF can form particles, one gigaton of allochthonous carbon, including microbes and proteins, can be bound and precipitated by fucoidan. However, further studies are required to constrain the resulting export flux and its potential attenuation, for example by quantifying the fucoidan carbon content in sinking particles obtained with sediment traps (**Fig. 4D**).

Our results suggest that the 0.5-1% of polymeric fucose carbon (0.5-1 µM FC) in the global ∼650 Gt dissolved organic carbon pool^10^ at least partially stems from ochrophytes. Universal presence of fucoidan exudation suggests it is an ancestral trait that evolved before the emergence of multicellularity in this phylum over 400 million years ago^51^. Without precipitation an annual release of ∼ 50 Mt soluble fucoidan would have accumulated and transitioned the ocean volume into a gel-state long time ago. Fortunately, precipitation removes fucoidan together with other molecules from solution, keeps their concentration low, and the ocean water clear and liquid. We conclude, fucoidan can act as a molecular key that ignites the biological carbon pump in the sunlit ocean.

## Methods summary

The accumulation and removal of dissolved organic molecules including the complex polysaccharide fucoidan was investigated along transects perpendicular to near-shore brown algae populations as well as in mesocosm experiments across the globe with brown macroalgae (**Fig. 1**). Six mesocosm experiments with specimens from either Laminariales (four) or Fucales (two) were conducted over 24 days, followed by year-long monitoring of incubation water (**Fig. S3A**). For the first 12 days, macroalgae specimens attached to their original substratum were kept in mesocosms next to control mesocosms without macroalgae (**Fig. 1A-D, F**). 1 L of mesocosm water was sampled every second day, followed by an exchange of half the mesocosm water with fresh seawater, while recording environmental conditions as well as primary productivity. After 12 days, macroalgae were taken out and sampling continued for another 12 days without water exchange. After 24 days of incubation, long-term incubations featuring quarter-yearly sampling started with 6-10 L of water from each mesocosm: one set up under a regulated light cycle at 20°C, another set up in darkness at 4°C with added inorganic nutrients corresponding to 40 µM NO_3_^−^ and 3 µM PO_4_^3-^, mimicking surface and deep ocean environmental conditions, respectively, not considering differences in pressure.

The analysis of obtained water and particulate samples consisted of specific quantification of an organic carbon or carbohydrate fraction complemented by visual analyses of stained cells and carbohydrates (**Fig. S3B-C**). Dissolved organic carbon was derived from total organic carbon measurements of filtered (pre-combusted glass fiber-filters 0.7 µm, GF/F) water. Polysaccharide concentrations and compositions of total dissolved fucoidan, surface active fucoidan and particulate fractions were determined. For total dissolved fucoidan quantification, GF/F filtered water was dialyzed using a 1 kDa membrane and retained molecules were hydrolyzed with 1 M hydrochloric acid at 100 degrees C for 24 h. Released monosaccharides were quantified by liquid chromatography with pulsed amperometric detection. The total dissolved fucoidan quantification method is faster and provides a stronger signal, as it also detects fucoidan molecules with lower charges including oligomers. For surface-active fucoidan quantification, an additional anion exchange chromatography step of glass fiber-filtered water was included, and the 4 M sodium chloride eluate was processed further via dialysis and acid hydrolysis. The AEX-based SAF quantification selectively concentrates fucoidan with high sulfate group density along the macromolecule^37,45^. The experimentally determined carbon content of fucoidan ranges between 25 and 31%, hence we used the factor 0.27 to convert fucoidan to fucoidan carbon^37^.

Fucoidan extracted from incubated specimens served as standards and was processed along with samples for both approaches. For particulate fucoidan, material on glass fiber filter pieces was hydrolyzed with 1 M hydrochloric acid and released monosaccharides quantified. Intact polysaccharides were detected in dialyzed water and on filters by probing with structure-sensitive monoclonal antibodies. Microbial cells were fixed on polycarbonate filters and counted after DAPI staining. A separate incubation of fucoidan and proteins in roller tanks was conducted to investigate their precipitation potential followed by flow cytometry analysis. A detailed description of the materials and experimental procedures can be found in the supplementary information.

## Acknowledgments

The authors thank technicians at Max-Planck Institute for Marine Microbiology and Marum Centre for Environmental Research, University of Bremen, namely Tina Horstmann for carbohydrate microarray analysis, Alek Bolte and Katharina Föll for HPAEC-PAD measurements and Gabriele Klockgether for particulate organic carbon measurements, as well as Fernanda Vargas Acevedo, Ricardo Calderón from Estación Costera de Investigaciones Marinas, Chile, Alistair Busby, Chris JT Boothroyd, Derek Kemp, John Bolton and Andrea Plos from the Department of Forestry, Fisheries and the Environment and the Department of Biological Sciences, University of Cape Town, South Africa, Lynne Butler, Bonny Clarke, the chief engineer, the captain and the crew of the R/V *Endeavor* during cruise EN683, Ariane Brandenburg and Chris Blake with the University of Waikato at Sulphur Point, Tauranga, New Zealand, M. Guadalupe Barba-Santos and Evelyn Raquel Salas-Acosta in the reef systems research unit at Puerto Morelos, Universidad Nacional Autónoma de México, Lena Eggers, Katherine Jones and Philipp Hach with the Alfred-Wegener Institute in Bremerhaven, Germany, the crew of R/V *Neomysis* and the Roscoff Aquarium Services platform part of EMBRC-France supported by the Investments of the Future program (ANR-10-INSB-02), for its technical assistance and support at the Station Biologique de Roscoff, France, and Joanna Norkko, Jaana Koistinen, Eva Rohlfer from Tvärminne Zoological Station for their assistance in field and lab work, and Phillip McFaul from Islander Kelp on Rathlin Island, Northern Ireland for support with water sampling inside their commercial kelp farm. The authors thank Hussein Awwad, Mohammad Shahin, Carla Knappik and Daria Gulenko for their support with sample processing. The authors thank Kai-Uwe Hinrichs, Roman Stocker and Bryce Inman for comments on the manuscript.

## Funding

The authors acknowledge invaluable support from the DFG Heisenberg grant for Glyco-Carbon Cycling in the Ocean and the Cluster of Excellence initiative (EXC-2077–390741603), Simons Collaboration on Principles of Microbial Ecosystems, ANID-Millennium Science Initiative Program – INCN2019-015 and NCN2023_004, ANID - FONDECYT (#1241901 and #1181719 to E.W), European Research Council grant for project C-Quest, the sea4SoCiety (03F0896), CDRmare campaign in the German Marine Research Alliance (for funding incubations in Finland and Sargassum sampling in Mexico), the Entrepreneurial Universities Macroalgal Biotechnologies Programme supported by the Tertiary Education Commission (TEC) and the University of Waikato, New Zealand, the Centre for Coastal Ecosystem and Climate Change Research (CoastClim) supported by the Jane and Aatos Erkko Foundation, the Ella and Georg Ehrnrooth Foundation, and the Research Council of Finland (# 354454 to CG), Finland, and funding from the U.S. National Science Foundation (OCE-1736772 and OCE-2022952 to CA) and the Max-Planck-Society.

## Author contributions

Conceptualization: I.H., H.B.-W. and J.-H.H. with contributions from all authors; methodology: I.H., H.B.-W. and J.-H.H.; investigation: I.H., H.B.-W., B.S.-M., E.W., M.K., M.R., C.A., C.L., S.G., D.P., P.P., T.T., M.M., C.G., L.W.; data curation: I.H., H.B.-W., T.T., D.B., M.S., C.U., L.W.; data analysis: I.H., H.B.-W., M.B., J.-H.H.; visualization: I.H., H.B.- W.; funding acquisition: J.-H.H.; M.M.; C.A.; writing – original draft: I.H., H.B.-W. and J.-H.H.; writing – review & editing: all authors.

## Declaration of interest

H.B.-W., J.-H.H., M.F.-M, M.P. are part of the advisory board of Seafields Solutions. M.F.-M. is founder of MacroCarbon. J.-H.H. and H.B.-W. have filed a patent on the quantification of fucoidan as a tracer for carbon sequestration and carbon credit enumeration by brown algae.

## Additional information

Supplementary Information is available for this paper.

## Data and materials availability

All data is available at PANGAEA^52^ data repository: https://doi.pangaea.de/10.1594/PANGAEA.980715,

https://doi.pangaea.de/10.1594/PANGAEA.980716

https://doi.pangaea.de/10.1594/PANGAEA.980717,

https://doi.pangaea.de/10.1594/PANGAEA.980718,

https://doi.pangaea.de/10.1594/PANGAEA.980719,

https://doi.pangaea.de/10.1594/PANGAEA.980720,

https://doi.pangaea.de/10.1594/PANGAEA.980721,

https://doi.pangaea.de/10.1594/PANGAEA.980722,

https://doi.pangaea.de/10.1594/PANGAEA.980723,

https://doi.pangaea.de/10.1594/PANGAEA.980724,

https://doi.pangaea.de/10.1594/PANGAEA.980725.

