## Supplementary information for "Macroalgal fucoidan can activate the biological carbon pump"

**The PDF file includes:**

Extended Materials and Methods
Code and intermediate results of hierarchical Bayesian models
Figs. S1 to S22
Tables S1 to S2
References (53–65)

### Extended Materials and Methods

#### *Experimental set-up*

Algae incubations were carried out at six locations across the world with locally abundant species of the order Laminariales or Fucales (**Table S1, 2**). *Lessonia trabeculata, Ecklonia maxima, Sargassum fluitans, Ecklonia radiata, Laminaria digitata* and *Fucus vesiculosus* were incubated in local ambient seawater (**Fig. S2**). Incubation volumes ranged between 40 and 250 L. Seawater was pre-filtered depending on water supply installed on the incubation systems used. For *L. trabeculata* and *E. maxima* incubations water was pre-filtered at 80 µm. *E. radiata* incubations started with unfiltered seawater and were topped up from day two onwards with seawater pre-filtered at 10 µm and UV treated, except on day ten when new, unfiltered seawater was used. For *S. fluitans*, *L. digitata* and *F. vesiculosus* incubations, unfiltered ambient seawater was used. *L. trabeculata*, *S. fluitans, E. radiata* and *L. digitata* were incubated outdoors under ambient light conditions, in the case of *L. trabeculata* and *E. radiata* under a transparent roof and 50% shade cloth respectively, whereas *E. maxima* and *F. vesiculosus* incubations were conducted indoors under artificial light conditions. Each mesocosm experiment consisted of six individual tanks, supplied with air stones for oxygen and an aquarium pump for water circulation. Water temperatures emulated local natural conditions ± 2°C by placing the incubation tanks of *L. trabeculata* (17°C), *E. maxima* (18°C) and *E. radiata* (18°C) into a constantly running and cooling water bath. *L. digitata* incubations were cooled to around 17°C using chillers (TECO TK500 G2, France), and *S. fluitans* incubations during the R/V *Endeavor* cruise EN683 were heated to 23°C using a heating rod (Eheim Thermocontrol 300, Germany) in the surrounding water bath, with a higher temperature deviation down to 16°C around day 8 due to a storm. *F. vesiculosus* was incubated at 14°C in a temperature-controlled room.

**Algae biomass** Total wet weight of individual specimens was quantified after the 12-day incubation. In addition, algae were separated into holdfast, stipe, and fronds and weighed independently. Biomass of *S. fluitans* in the Sargasso Sea experiment was freeze dried, whereas all the other algae were dried at 60°C in an oven to determine dry biomass.

**Primary productivity** Primary productivity was measured using PyroScience FireSting oxygen probes (Pyroscience GmbH, Germany) during all incubations, except for *L. trabeculata* incubations, where no measurements were taken. Measurements were taken at the beginning, after 2 to 4 days and at the end of the incubation. For each species and replicate, a single algal frond was incubated for a minimum of 20 min inside a separate 2 L bottle filled with tank water and measuring probes during day and night. In the case of *S. fluitans* mesocosms, whole specimens were fit into the 2 L bottle. The water was stirred using an underwater propeller (Tamiya, Japan). Temperature and oxygen were measured. Based on oxygen production and respiration, we calculated the net primary productivity in µmol $O_2$ per dry biomass per day (based on a 1:1 conversion factor). This data was used to calculate the fixed carbon and the ratio of % NPP in mg fucoidan carbon $g^{-1}$ $d^{-1}$ to mg fixed carbon $g^{-1}$ $d^{-1}$.

**PAR measurements** PAR data for the *L. trabeculata* incubation was obtained from the website of the Centro de Estudios Avanzados en Zonas Áridas (CEAZA) approximated based on a study [53]. Mesocosms in the *E. maxima* experiment were each irradiated with an Osram Power Star hqi-e 440W/D lamp located centrally above mesocosms with a nominal flux of 40000 lumen for 14 hours daily. The PAR data during the *S. fluitans* incubation was recorded by the R/V *Endeavor*.

Estimates for PAR during the *L. digitata* incubation experiment at Roscoff, France, were calculated from NASA POWER (https://power.larc.nasa.gov/). PAR measurements were conducted using a HOBO logger (HOBO MX2202, Onset, US) during *E. radiata* incubations and using an RBRsolo logger with an attached PAR sensor (LI-192 Underwater Quantum Sensor) during *F. vesiculosus* incubations. Each mesocosm with *F. vesiculosus* was irradiated with AQUAMEDIC Aquarius 120 LED lights and set to a 12/12 light cycle, including two hours of dusk and dawn. The PAR loggers were placed inside the filled mesocosms at the level of the incubated algae, whereas for the other incubations light was measured in air.

**Transects** Water samples were taken along transects moving away from seaweed beds at all local mesocosm sites. In Chile, two transects were sampled starting from Algarrobo and Las Cruces. Niskin-bottle sampling was performed at 15 m depth at station 1, 5 m and 30 m depth at station 2, 5 m and 60 m depth at station 3, and 5 m and 90 m depth at station 4. In South Africa, station 1 was sampled at 15 m depth, station 2 at 5 m and 30 m depth, station 3 at 5 m and 45 m depth, and station 4 at 5 m and 60 m depth. On cruise R/V *Endeavor* EN683, sampling was conducted at three stations, station 21 to 23, at 5 m depth, the deep chlorophyll maximum (DCM), minimum dissolved oxygen ($O_2$min), and the bottom depth. In Roscoff, France, samples were collected during incoming and outgoing tides at four distinct stations at 5m and 30 m depth. In New Zealand, surface and deep samples were taken at four stations. In Finland surface water samples were sampled at five stations moving away from *F. vesiculosus* beds. Additionally, sampling included inside and outside measurements of *Sargassum* patches in Mexico. Sampling was conducted at a *Saccharina latissima* farm in Northern Ireland upstream, inside and downstream of the algae farm at 5 m depth. At each station three replicates from the same Niskin bottle cast were analyzed. Detailed information on sampling locations can be found in supplementary **Table S1**.

***Sample analysis***

Every second day for the first 24 days of incubation, 1 L of water was sampled from each tank (**Fig. S3B**). Afterwards, every 3 months, 1 L of water was taken from the long-term incubations. 800 mL of the sample was filtered through a pre-combusted (450°C, 4.5 h) 47 mm glass fiber filter (0.7 µm pore size). 20 mL of filtered water were acidified for analysis of dissolved organic carbon. Anionic polysaccharides were extracted and purified from 400 mL of the filtered seawater using anion exchange chromatography (AEX) and eluted with 4 M sodium chloride (NaCl). 7.5 mL of unfiltered water for bacterial cell numbers was fixed in a final concentration of 3% formaldehyde solution (v/v) for 1 h at room temperature. The solution was filtered onto a 47 mm polycarbonate filter with a pore size of 0.22 µm, dried and frozen at -20°C. After 24-days of incubation, tank water was separated into long-term incubations before emptying tanks. Sedimented particles were scooped out from the bottom of the tanks in Chile, South Africa and France (**Fig. S3C**). Particles were frozen at -20°C until further analysis. Samples were transported frozen to the Max Planck Institute for Marine Microbiology in Bremen, Germany, for further purifications, monosaccharide quantification, antibody binding and cell counting.

**DOC** Dissolved organic carbon (DOC) was measured at Tvärminne Zoological Station, Hanko, Finland for *F. vesiculosus* incubations. Samples were acidified to pH 2 using hydrochloric acid (HCl, 25%, p.a., Carl Roth, Germany) in pre-combusted glass vials with acid-washed lids. All samples were stored at 4°C in the dark until analysis. For all other incubations, DOC was measured at the Institute for Chemistry and Biology of the Marine Environment, University of Oldenburg, Germany. Filtered samples were frozen and stored at -20°C and acidified prior to measurements in glass vials with acid-washed lids. Concentrations of DOC were determined using a Shimadzu

TOC-$V_{CPH}$-analyzer. DOC deep sea reference material (DSR; Hansell Biogeochemistry Laboratory, University of Miami, USA) was used for quality control, with accuracy and precision < 5%.

**Dialysis** To desalt and size select samples, 10 mL of filtered seawater or AEX fractions were transferred to pre-rinsed 1-kDa membranes (SpectraPor® Biotech CE Dialysis Membrane, Carl Roth, Germany) and dialyzed three times against MilliQ-water at 4°C for a minimum of 12 hours in total. Dialyzed samples were transferred to 15 mL tubes, freeze dried (Labogene, Scanvac Coolsafe, Denmark) and resuspended in 2 mL MilliQ water.

**Acid hydrolysis** To convert polysaccharides into quantifiable monosaccharides, 500 µL of the dialyzed, resuspended samples were combined with 500 µL 2 M HCl in pre-combusted glass vials (450°C, 4.5 h), which were sealed. Polysaccharides were hydrolyzed at 100°C for 24 h. After hydrolysis, the samples were transferred to microtubes and dried in an acid-resistant vacuum concentrator (Martin Christ Gefriertrocknungsanlagen GmbH, Germany). Samples were resuspended in 500 µL MilliQ- water.

**Monosaccharide quantification** Monosaccharides were quantified using anion exchange chromatography with pulsed amperometric detection (HPAEC-PAD), as previously described[54,26]. Samples were analyzed using a Dionex ICS-5000+ system equipped with a CarboPac PA10 analytical column (2 x 250 mm) and a CarboPac PA10 guard column (2 x 50 mm). Neutral and amino sugars were separated during an isocratic phase with 18 mM sodium hydroxide (NaOH) followed by a gradient reaching up to 200 mM sodium acetate ($NaCH_3COO$) to resolve acidic monosaccharides.

**Carbohydrate microarray analysis** Monoclonal antibodies targeting brown algae polysaccharides were employed to screen mesocosm samples for dissolved carbohydrates. Filtered and dialyzed samples were concentrated by a factor of 20 prior to analysis with carbohydrate microarrays by printing aliquots onto nitrocellulose membrane and measuring antibody binding via colorimetric signal[26,55]. Extracted fucoidan from the incubated algal biomass, alginate (Sigma), laminarin (Carbosynth) and mixed-linked glucan (Megazyme) were used as standards at concentrations of 2.5, 5, 10 and 20 mg $L^{-1}$. The printed microarrays were blocked in milk phosphate buffered saline (MPBS, 5% skim milk in PBS) for 1 h and incubated individually for 2 h at a concentration of 10 µg $mL^{-1}$ with the monoclonal antibodies BAM1, BAM2, BAM7, LM21, BS400-2, BS400-4 and the anti-rat control [56,57]. The arrays were washed with phosphate buffered saline (PBS) and incubated with secondary antibodies in MPBS for 2 h, then washed again in PBS and deionised water. The arrays were developed in 5-bromo-4-chloro-3-indolyl phosphate and nitroblue tetrazolium in alkaline phosphatase buffer (100 mM NaCl, 5 mM MgCl2, 100 mM Tris-HCl, pH 9.5) and analyzed with the software Arrray-Pro Analyzer 6.3 (Media Cybernetics).

**Fucoidan extraction from seawater** Fucoidan was extracted from seawater using anion exchange columns for AEX adapted from [37]. 400 mL of GF/F filtered seawater samples passed through 5 mL HiTrap ANX high sub FF columns (Cytiva), which had been conditioned with 25 mL of 0.5 M NaCl and 20 mM Tris-HCl (pH 8), 25 mL of 4 M NaCl and 20 mM Tris-HCl (pH 8) and again with 25 mL of 0.5 M NaCl and 20 mM Tris-HCl (pH 8), all at 5 mL $min^{-1}$ flow rate. For washing, 30 mL of 0.5 M NaCl and 20 mM Tris-HCl (pH 8) was applied after the sample and combined with the sample flow through. To elute anionic fucoidan, 12 mL of 4 M NaCl and 20 mM Tris-HCl (pH 8) was passed over the columns at 5 mL $min^{-1}$ and collected in a 15-mL tube. The 15-mL tubes were stored at 4°C until further processing. The columns were cleaned with 25 ml MilliQ at 5 mL min$^{-1}$ flow rate, 20 mL 1 M NaOH at 1 mL min$^{-1}$ flow rate and 25 ml MilliQ at 5 mL min$^{-1}$ flow rate.

**Fucoidan extraction from biomass** Marcoalgal biomass was dried at 60°C in an oven or in case of *Sargassum* biomass freeze-dried. Dried biomass was processed and fucoidan extracted as recently described [58]. The obtained fucoidan powder was used to prepare fucoidan standards.

**Particulate organic carbon** GF/F filters were cut, with 20-35% of the total cut filters weighed and placed in an acid desiccator overnight. The filter pieces were dried in an oven at 60°C for 2 hours, packed into tin cups, and compressed. Particulate organic carbon (POC) was quantified using a Vario Micro Cube (Elementar Analysensysteme) calibrated with sulfanilamide standards.

**Particulate fucoidan** GF/F filters were cut and weighed. 15% of the cut GF/F filters were directly acid hydrolyzed in 600 µL of 1M HCl at 100°C for 24 h in glass vials. The supernatant was transferred, dried and resuspended in the same amount. Monosaccharides were quantified using HPAEC-PAD (see above).

**Sequential extraction and ELISA BAM1** From approximately 20% of each GF/F filter, polysaccharides were sequentially extracted using MilliQ-water and 0.3 M EDTA. Filter pieces were mixed with 1.8 mL of MilliQ-water, vortexed and kept in an ultrasonic water bath for 1 h. Filters were centrifuged at 6000 x g for 15 min. Supernatant was transferred into a new vial and filter pieces were mixed with 1.8 mL of 0.3 M EDTA, vortexed and kept in an ultrasonic water bath for 1 h. Filters were centrifuged at 6000 x g for 15 min and supernatant was again transferred into a new vial. Extracts of MilliQ and EDTA were combined for further analysis by pipetting 50 µl of each into a pre-coated 96 well plate to settle overnight at 4°C. Solution in each well was removed the next day, 200 µL of MPBS was added to each well and incubated for 2 h. After incubation, MPBS was removed and wells were washed 9 times with deionized water. 100 µL of antibody BAM1 in MPBS solution at a concentration of 1:10 was added into each well and incubated for 1.5 h. The antibody-mix was then removed and wells were washed 6 times with deionized water. 100 µL of secondary antibody anti-rat in MPBS solution at a concentration of 1:10000 was added into each well and incubated for 1.5 h. Antibody-mix was removed afterwards and wells were washed 6 times with deionized water. Development solution was added and stopped after 15 min with 1M HCl. Absorption was measured at 450 nm using Spectramax Id3 plate reader (Molecular Devices).

**Immunolabeling on filters** Frozen filters were cut into pieces and filter sections were incubated in 400 µl MPBS for 1 hour inside tubes. After blocking, the solution was replaced with 200 µl of primary antibody BAM1 at 1:5 dilution in MPBS for 1.5 hours. After the incubation, the filter sections were washed 4 times with 1 mL of PBS. 200 µl of secondary antibody anti-rat FITC conjugated at 1:100 dilution were added to filters inside the tube and incubated for 1.5 hours in darkness. Finally, the filter sections were washed 4 times with 1 mL of PBS and stained with DAPI (see below).

**Cell counts** Frozen filters were cut into pieces and filter sections were mounted on glass slides with premixed 4',6-Diamidino-2-phenylindole (DAPI) 1:1000 diluted in the antifading reagents Citifluor:Vectashield as 4:1 (v/v).

**Microscopy** Slides were covered and stored at 4°C in the dark. Cells were counted and immunolabeled filter sections for BAM1 fucoidan binding were visualized using a Nikon 50i microscope equipped with a Zeiss AxioCam MRc camera at 100 x magnification (Plan Apo VC objective). Sedimented particles were imaged using Thermo Scientific Invitrogen EVOS FL Auto Imaging System at 20x magnification Brightfield.

***Roller tank experiment***

A separate experiment was carried out to test the precipitation potential of fucoidan. Commercial crude fucoidan from *Laminaria hyperborea* was purified by using a sequence of precipitation with $CaCl_2$, followed by anion exchange chromatography[45], and a precipitation and desalting method using ethanol [59]. In brief, a 5 g/L fucoidan solution was treated with a 2 M $CaCl_2$ solution in an alkaline environment created with NaOH. The solution was centrifuged to separate precipitated impurities from dissolved fucoidan in the supernatant. For the subsequent anion exchange chromatography, the fucoidan solution was neutralized and diluted in running buffer A (0.5 M NaCl, 20 mM Tris-HCl) and loaded on an ANX-HiTrap column. After washing with buffer A the purified anionic polysaccharide was eluted with running buffer B (4 M NaCl, 20 mM Tris-HCl) in 12 mL per 1 mg crude fucoidan. To desalt the samples, extracted fucoidan in the eluants was precipitated using 70-80% EtOH, filtered over a combusted (450°C, 4.5h) GF/F filter and washed with 70-80% EtOH. The white solid was reconstituted from the filter in ultra-pure water and in the final step freeze dried. A 5 mg/mL pure fucoidan solution was cooled in an ice bath and the pH was adjusted to 8. A DAMP (2.5 M) solution was added to buffer the system during the reaction. With addition of a CDAP solution (100 mg/mL in ACN) the reaction was started and the pH value was monitored over 15 min. Afterwards the reaction mixture was directly loaded on a self-packed SEPHADEX G-50 column with a borate buffer (0.2 M) as mobile phase. For the column chromatography an AKTA Start system was used according to [60]. The eluted activated sugar was collected, and 4 mg of fluorescein amine were added. The solution was allowed to react overnight and subsequently purified with another column chromatography using a SEPHADEX G-50 based column and a phosphate buffer (50 mM, 100 mM NaCl) as mobile phase. The solvent of the eluted fluorescently labelled fucoidan was exchanged by ultra-pure water with ultrafiltration tubes (MW cut off at 5 kDa).

To this end, the extracted fucoidan was fluorescently labeled and incubated in 3.5% artificial seawater (Sigma) at final concentrations of 0.01, 0.05, and 0.1 mg $L^{-1}$. In addition, triplicate incubations of fucoidan at these concentrations in combination with the protein lysozyme at a fucoidan:protein mass ratio of 1:100, corresponding to lysozyme concentrations of 1, 5, and 10 mg $L^{-1}$, were set up. Lysozyme-only treatments at the same concentrations (1, 5, and 10 mg $L^{-1}$) were included as protein controls. Artificial seawater without added fucoidan or protein served as a blank control. Samples of 12.5 mL were incubated in 15mL-falcon tubes, wrapped in aluminum foil to keep them dark at 16°C on a rolling table at 10 rpm to ensure continuous gentle mixing. Incubations were carried out for a total duration of 16 days. Subsamples of 500 µL were collected on days 0, 2, 5, 8, 13, and 16. Immediately after collection, samples were stored at 4°C for analysis once the incubation period concluded.

All samples were analyzed using flow cytometry (BD Biosciences FACSCalibur™). Forward scatter (FSC), side scatter (SSC), and fluorescence signals were recorded. The detection thresholds for side scatter (SSC) and fluorescence channel FLH1 were set to zero, allowing detection of all particles within the size detection range of the flow cytometer, regardless of fluorescence labeling. This approach ensured inclusion of both fluorescent and non-fluorescent particles in the analysis. Particles were counted from the signature plot of SSC-H vs green fluorescence. The output was analysed using the BD CellQuest™ Pro software v5.2.1.

***Data analysis***

Fucoidan concentrations were calculated from fucose HPAEC-PAD data based on fucose concentrations in acid-hydrolyzed fucoidan standards. Fucoidan was extracted from algae biomass and added to seawater at different concentrations. The dissolved fucoidan in seawater was then processed with mesocosm and transect samples. Fucoidan standards were added to GF/F filter pieces and processed together with samples. In this way, specific fucoidan standard series were generated for samples before and after charge-selection and particulate samples for each algal species. The calibrated dissolved fucoidan concentrations served to calculate a cumulative fucoidan value by accounting for water exchanges during the incubation of brown algae. For each independent mesocosm until day 12, half the concentration measured during the previous sampling prior to the water exchange was added to calculate the cumulative value. The cumulative values were used to calculate the change in fucoidan concentrations between two sampling events. As this value was highly susceptible to measurement error, we also calculated a sliding window-change in fucoidan concentrations which spanned four sampling events. All values were calculated separately for total dissolved fucoidan and the surface-active fucoidan (SAF) fraction selected based on size and charge.

To assess the drivers of fucoidan accumulation and removal in mesocosms, a series of Bayesian hierarchical models was computed using the package Rstantools in R version 4.3.3[61,62] (see supplementary section **Bayesian Models**). Mesocosm incubations with *E. maxima* (Cape Town, South Africa) and *L. digitata* (Roscoff, France) were excluded from the analysis due to light limitation in the case of *E. maxima* (**Fig. S5, S22**) and missing water exchanges in the case of *L. digitata* (**Fig. S4**). First, two models that included either the actual concentrations of dissolved fucoidan and the SAF fraction or the cumulative concentrations as dependent variables were constructed, with presence/absence of brown algae and incubation period as fixed factors and location and day of incubation as random factors. Fucoidan concentrations were assumed to be normally distributed. Second, two models that used the two-point or the sliding window-changes in dissolved fucoidan and the SAF fraction concentrations dependent on brown algae biomass scaled to mesocosm volume and fucoidan concentration during the previous sampling event were computed. Changes were assumed to be normally distributed.

The posterior distributions of the effect of brown algae biomass and previous fucoidan concentration were validated by simulating mesocosm incubations (**Fig. S21)**. Random effects of location and mesocosm were included in the simulation. As during the actual experiments, water exchanges were simulated to account for the diluting effect on fucoidan concentrations. Biomass per volume ratios were drawn from the physical mesocosm experiments. These posterior distributions were used in Monte Carlo simulations to project the global release of fucoidan. The simulations used published net primary productivity per area data with assumed gaussian distribution (mean = 0.546 kg C $m^{-2}$ $yr^{-1}$, std. dev. = 0.632 kg C $m^{-2}$ $yr^{-1}$, n = 142) and the global subtidal brown algae area with assumed gaussian distribution (mean = 1.68 x $10^{12}$ $m^2$, 25$^{th}$ percentile = 1.43 x $10^{12}$ $m^2$, 75$^{th}$ percentile = 1.79 x $10^{12}$ $m^2$, [50], together with the linear model correlating net primary production and frond biomass for the studied algal species (**Fig. 2A**), and the Bayesian posterior distributions of the effect of brown algal biomass on fucoidan accumulation. One thousand runs were performed for each total dissolved fucoidan and the SAF fraction.

Persistence of fucoidan concentrations in all incubations combined was tested using paired statistical tests. Normality was tested using Shapiro-Wilk test. Differences between mesocosms with brown algae and mesocosms without brown algae were tested for normally distributed data with paired t-tests. Due to predominantly non-normally distributed data, Wilcoxon tests were employed (**Fig. S11**). Spearman's correlation of particulate organic carbon and particulate fucoidan concentrations over time for all incubations combined were determined, fitted lines were modelled with linear regression.

Literature values on fucose content in marine particulate organic matter was collected by performing a google scholar search on March 28, 2025 with the search strings “particulate”, “carbohydrate”, “composition” and “fucose”. Articles on pages 1 through 30 were included in the data. Information on particulate organic carbon and particulate fucose was extracted. To estimate the contribution of fucoidan C/POC, data was $\log_{10}$-transformed and linear models were calculated, excluding measurements taken in riverine and estuary environments (**Fig. 4A**).

The occurrence map (**Fig. 1**) was plotted using R packages ‘marmap’[63] and ‘sf’[64].

**Bayesian models** Code and intermediate results show the hierarchical Bayesian models used to analyze the fucoidan data. The models are fitted to dissolved fucoidan and SAF concentrations, cumulative fucoidan concentrations, and the change in fucoidan concentration over time (dC/dt). The models predict the effects of treatment (with or without brown algae), incubation period (day 0-12: synthesis; day 14-24: degradation), and random effects of location and sampling day. As conservative approach, the models use uniform distributions as uninformative priors.

First, load required packages, set working directory and import data.

```
dat4saving = read.csv("20250625_fucoidanBCP_data.csv")
```

***fit1: Comparison of fucoidan estimates over incubation period***

This hierarchical Bayesian model evaluates the impact of treatment and incubation period on dissolved fucoidan and SAF estimates. Location and sampling day are included as random factors. To conserve computing time, a saved rds file with the model results is loaded.

```
# load fit1 or rerun

#fit1 <- brm(
#  bf(Fucoidan_mgL_dissolved ~ Treatment * Period + (1 | Location / Day))+
#    bf(Fucoidan_mgL_SAF ~ Treatment * Period + (1 | Location / Day)),
#  data = dat4saving, family = gaussian(), chains = 4, cores = 2, iter = 2000,
#  control = list(adapt_delta = 0.95))
# save fit1 to file
#saveRDS(fit1, file = "20250620_model_fit1.rds")

fit1 <- readRDS("20250620_model_fit1.rds")

#Summary of the model fit2 results
broom.mixed::tidy(fit1) |>
  select(response, term, estimate, std.error, conf.low, conf.high)

## # A tibble: 14 × 6
##    response              term                  estimate std.error conf.low conf.high
##    <chr>                 <chr>                    <dbl>     <dbl>    <dbl>     <dbl>
##  1 FucoidanmgLdissolved (Intercept)              1.19      0.487    0.211     2.19
##  2 FucoidanmgLSAF       (Intercept)              0.219     0.0798   0.0545    0.367
##  3 FucoidanmgLdissolved TreatmentControl        -0.564     0.139   -0.838    -0.288
##  4 FucoidanmgLdissolved Periodsynthesis         -0.297     0.146   -0.589    -0.00861
##  5 FucoidanmgLdissolved TreatmentControl:…       0.0585    0.183   -0.315     0.402
##  6 FucoidanmgLSAF       TreatmentControl        -0.128     0.0289  -0.183    -0.0707
##  7 FucoidanmgLSAF       Periodsynthesis          0.0309    0.0297  -0.0269    0.0892
##  8 FucoidanmgLSAF       TreatmentControl:…      -0.0491    0.0386  -0.126     0.0250
```

```
##  9 Location             sd__NA.Fucoidanmg…   1.10     0.438    0.548     2.21
## 10 Location:Day         sd__NA.Fucoidanmg…   0.283    0.0771   0.109     0.426
## 11 Location             sd__NA.Fucoidanmg…   0.168    0.0848   0.0798    0.373
## 12 Location:Day         sd__NA.Fucoidanmg…   0.0449   0.0188   0.00386   0.0788
## 13 FucoidanmgLdissolved sd__Observation      0.815    0.0360   0.749     0.888
## 14 FucoidanmgLSAF       sd__Observation      0.172    0.00753  0.158     0.188

# A Bayesian analog to the frequentist's R2 for fit1
print(bayes_R2(fit1))

##                       Estimate  Est.Error      Q2.5     Q97.5
## R2FucoidanmgLdissolved 0.5192822 0.03225382 0.4499583 0.5753442
## R2FucoidanmgLSAF       0.4192960 0.03827215 0.3430876 0.4903921
```

Next, the posterior predictive distribution of the model fit1 is obtained.

```
new_data = data.frame(Location = character(),
                      Tank = numeric(),
                      Day = numeric(),
                      Treatment = character(),
                      Period = character())
for (loc in unique(dat4saving$Location)){
  for (tank in unique(dat4saving$Tank)){
    for (treat in c("Algae", "Control")){
      for (period in c("synthesis", "degradation")){
        day = runif(1, 0, 24)
        new_data = rbind(new_data, c(loc, tank, day, treat, period))
      }
    }

  }
}
colnames(new_data) = c("Location", "Tank", "Day", "Treatment", "Period")
fit1_p.epred = fit1 |>
  linpred_draws(newdata = new_data, ndraws = 5, allow_new_levels = T)
fit1_p.epred$Period = factor(fit1_p.epred$Period, levels = c("synthesis", "degradation"))
strip_fills =  strip_themed(background_x = elem_list_rect(fill = pal_types))
strip_labels = c(FucoidanmgLdissolved = "dissolved fucoidan",
                 FucoidanmgLSAF = "SAF")
```

The posterior predictive distribution of the model fit1 is plotted, excluding due to experimental issues *Ecklonia maxima* at location South Africa (SA) and *Laminaria digitata* at location Roscoff (RO) in France, which are separately plotted posterior predictions (cross hairs and squares) and Fig. S22.

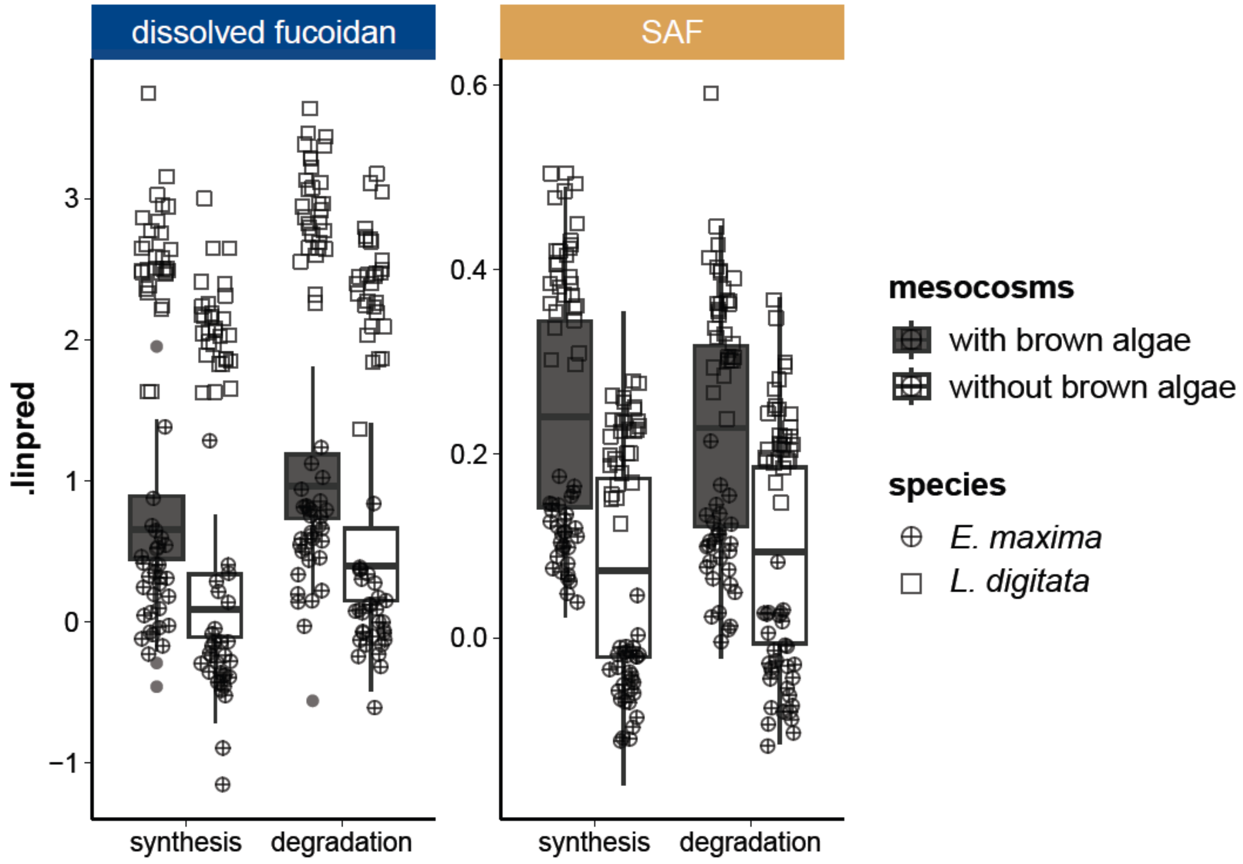


***fit2: Comparison of cumulative fucoidan estimates taking water exchanges into account***

This hierarchical Bayesian model has the same structure as the model fit1, but it uses as response variables the cumulative concentrations of dissolved fucoidan and SAF, which were calculated to account for water exchanges during the first 12 days of incubations and are contained in the data (dat4saving). To conserve computing time, a saved rds file with the model results is loaded.

```
# load fit2 or rerun
#fit2 <- brm(
#  bf(Cumulative_fucoidan_mgL_dissolved ~ Treatment * Period + (1 | Location/Day))+
#    bf(Cumulative_fucoidan_mgL_SAF ~ Treatment * Period + (1 | Location/Day)),
#  data = dat4saving, family = gaussian(), chains = 4, cores = 2, iter = 2000,
#  control = list(adapt_delta = 0.9)
# )
# save fit1 to file
#saveRDS(fit2, file = "20250620_model_fit2.rds")
fit2 = readRDS("20250620_model_fit2.rds")

# Summary of the model fit2 results
broom.mixed::tidy(fit2) |>
  select(response, term, estimate, std.error, conf.low, conf.high)

## # A tibble: 14 × 6
##    response                        term     estimate std.error conf.low conf.high
##    <chr>                           <chr>       <dbl>     <dbl>    <dbl>     <dbl>
##  1 Cumulativefucoidanmgldissolved  (Interc…    1.72      0.520    0.674      2.80
##  2 CumulativefucoidanmgLSAF        (Interc…    0.467     0.121    0.215      0.708
```

```
## 3 CumulativefucoidanmgLdissolved Treatme… -0.856 0.172 -1.20 -0.524
## 4 CumulativefucoidanmgLdissolved Periods… -0.276 0.161 -0.591 0.0424
## 5 CumulativefucoidanmgLdissolved Treatme… -0.0465 0.217 -0.466 0.389
## 6 CumulativefucoidanmgLSAF Treatme… -0.330 0.0395 -0.408 -0.254
## 7 CumulativefucoidanmgLSAF Periods… -0.0715 0.0430 -0.157 0.0132
## 8 CumulativefucoidanmgLSAF Treatme… 0.0334 0.0504 -0.0641 0.133
## 9 Location sd__NA.… 1.21 0.501 0.610 2.56
## 10 Location:Day sd__NA.… 0.0974 0.0704 0.00369 0.260
## 11 Location sd__NA.… 0.267 0.124 0.128 0.595
## 12 Location:Day sd__NA.… 0.0905 0.0199 0.0505 0.130
## 13 CumulativefucoidanmgLdissolved sd__Obs… 0.921 0.0375 0.852 1.00
## 14 CumulativefucoidanmgLSAF sd__Obs… 0.208 0.00976 0.190 0.228
```

```
# R2 for fit2
print(bayes_R2(fit2))
```

```
## Estimate Est.Error Q2.5 Q97.5
## R2CumulativefucoidanmgLdissolved 0.4658067 0.03171594 0.3996101 0.5232469
## R2CumulativefucoidanmgLSAF 0.6158605 0.02757619 0.5569149 0.6650069
```

***Rerun the model fit2 excluding locations SA and RO due to experimental issues.***

```
# load fit2_woSARO or rerun

#fit2_woSARO <- brm(
# bf(Cumulative_fucoidan_mgL_dissolved ~ Treatment * Period + (1 | Location/Day))+
# bf(Cumulative_fucoidan_mgL_SAF ~ Treatment * Period + (1 | Location/Day)),
# data = dat4saving[which(!dat4saving$Location %in% c("SA", "RO")),],
# family = gaussian(), chains = 4, cores = 2, #iter = 2000,
# control = list(adapt_delta = 0.99)
#)
#saveRDS(fit2_woSARO, file = "20250620_model_fit2_woSARO.rds")

fit2_woSARO = readRDS("20250620_model_fit2_woSARO.rds")

print("Summary of the model fit2_woSARO results:")
```

```
## [1] "Summary of the model fit2_woSARO results:"
```

```
broom.mixed::tidy(fit2_woSARO) |>
  select(response, term, estimate, std.error, conf.low, conf.high)
```

```
## # A tibble: 14 × 6
## response term estimate std.error conf.low conf.high
## <chr> <chr> <dbl> <dbl> <dbl> <dbl>
## 1 CumulativefucoidanmgLdissolved (Interc… 1.64 0.464 0.708 2.58
## 2 CumulativefucoidanmgLSAF (Interc… 0.544 0.217 0.100 0.989
## 3 CumulativefucoidanmgLdissolved Treatme… -1.14 0.181 -1.51 -0.787
## 4 CumulativefucoidanmgLdissolved Periods… -0.372 0.166 -0.701 -0.0368
## 5 CumulativefucoidanmgLdissolved Treatme… 0.172 0.235 -0.286 0.631
## 6 CumulativefucoidanmgLSAF Treatme… -0.424 0.0482 -0.520 -0.329
## 7 CumulativefucoidanmgLSAF Periods… -0.102 0.0545 -0.210 0.00381
## 8 CumulativefucoidanmgLSAF Treatme… 0.0774 0.0616 -0.0416 0.197
## 9 Location sd__NA.… 0.779 0.533 0.263 2.22
```

```
## 10 Location:Day                 sd__NA.…   0.0966    0.0706  0.00322   0.263
## 11 Location                     sd__NA.…   0.388     0.312   0.128     1.21
## 12 Location:Day                 sd__NA.…   0.106     0.0251  0.0571    0.155
## 13 CumulativefucoidanmgLdissolved sd__Obs…   0.835     0.0421  0.757     0.923
## 14 CumulativefucoidanmgLSAF       sd__Obs…   0.216     0.0125  0.193     0.242
```

```
# R2 for fit2_woSARO
print(bayes_R2(fit2_woSARO))
```

```
##                                 Estimate  Est.Error      Q2.5     Q97.5
## R2CumulativefucoidanmgLdissolved 0.4069709 0.04057954 0.3204582 0.4800066
## R2CumulativefucoidanmgLSAF       0.6497993 0.03049240 0.5832423 0.7009459
```

Obtain the posterior predictive distribution of the model fit2_woSARO.

```
new_data = data.frame(Location = character(),
                      Tank = numeric(),
                      Day = numeric(),
                      Treatment = character(),
                      Period = character())
for (loc in unique(dat4saving$Location[which(!dat4saving$Location %in% c("SA", "RO"))])){
  for (tank in unique(dat4saving$Tank)){
    for (treat in c("Algae", "Control")){
      for (period in c("synthesis", "degradation")){
        day = runif(1, 0, 24)
        new_data = rbind(new_data, c(loc, tank, day, treat, period))
      }
    }

  }
}
colnames(new_data) = c("Location", "Tank", "Day", "Treatment", "Period")
fit2_p.epred = fit2_woSARO |>
  linpred_draws(newdata = new_data, ndraws = 5, allow_new_levels = T)
fit2_p.epred$Period = factor(fit2_p.epred$Period, levels = c("synthesis", "degradation"))
strip_fills =  strip_themed(background_x = elem_list_rect(fill = pal_types))
strip_labels = c(CumulativefucoidanmgLdissolved = "dissolved fucoidan",
                 CumulativefucoidanmgLSAF = "SAF")
```

Create boxplots of the posterior predictive distribution of the model fit2_woSARO, excluding locations SA and RO due to experimental issues. The boxplots show the predicted cumulative fucoidan concentrations for the different treatments and periods.

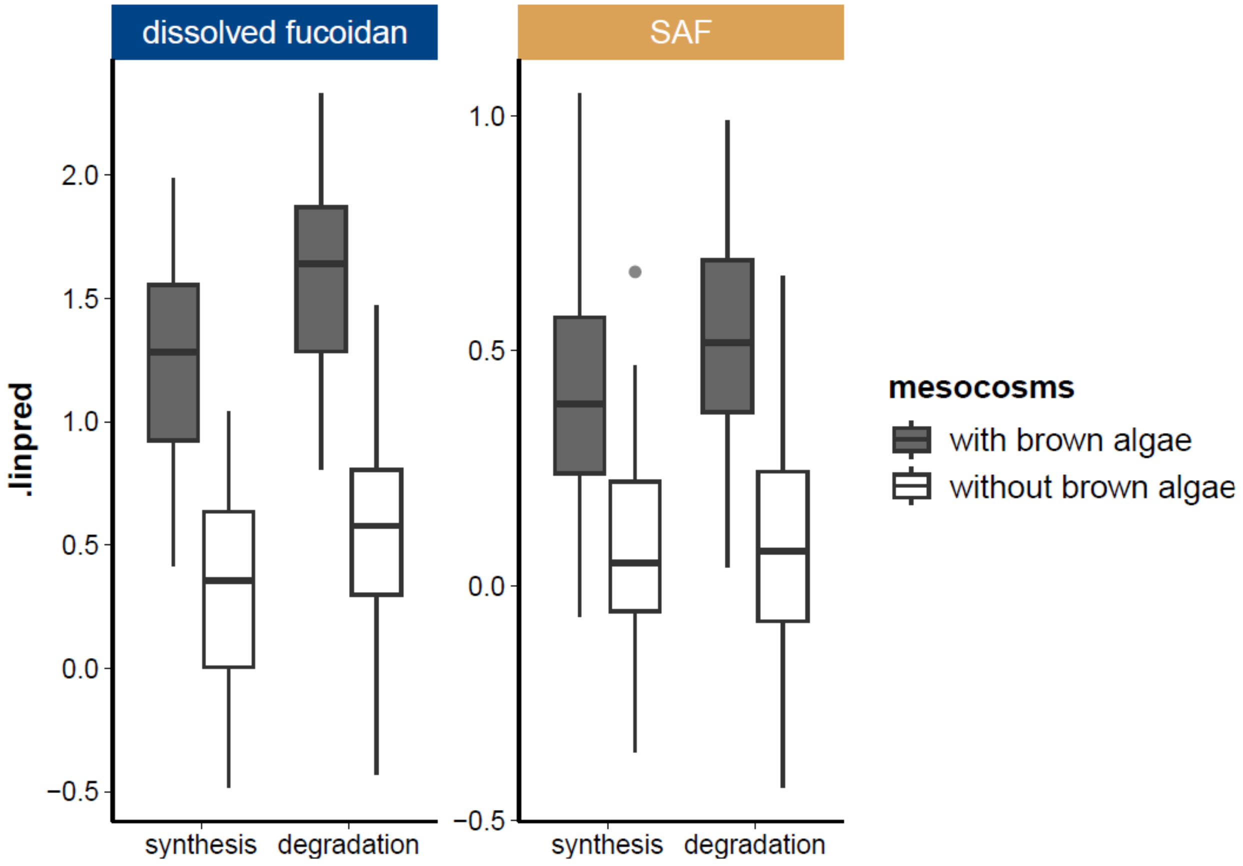


***fit3: Modeling the change in fucoidan concentration over time (dC/dt)***

The models fit1 and fit2 only coarsely distinguish between the “synthesis period” (days 0 - 12), after which brown algae were removed from mesocosms, and the “degradation period” (days 14 - 24). Therefore, assess the rate of change during the incubation, we fit a model that uses the change in fucoidan concentration over time (dC/dt) as response variable. This model is fitted to the change in cumulative concentration calculated as sliding window of 4 days to correct for technical noise. The model includes the brown algal biomass normalized to mesocosm volume (BMperV_gL) and the actual fucoidan concentration at the beginning of the sliding window as predictors. The model is fitted separately for dissolved fucoidan and SAF, excluding locations SA and RO due to experimental issues. To conserve computing time, a saved rds file with the model results is loaded.

```
# load model fit3_dissolved or rerun

#fit3_dissolved = brm(
#  bf(dCdt_sliding_mgLd_dissolved ~ BMperV_gL + Reference_fucoidan_mgL_dissolved +
#     (1 | Location/TankID)),
#  data = dat4saving[which(!dat4saving$Location %in% c("SA", "RO")),],
#  chains = 4, cores = 4, iter = 3000, warmup = 1000, thin = 2,
#  control = list(adapt_delta = 0.99))
#saveRDS(fit3_dissolved, file = "20250620_model_fit3_dissolved.rds")

fit3_dissolved = readRDS("20250620_model_fit3_dissolved.rds")
```

```
# Summary of the model fit3_dissolved results:
broom.mixed::tidy(fit3_dissolved)

## # A tibble: 6 × 8
##   effect   component group          term  estimate std.error conf.low conf.high
##   <chr>    <chr>     <chr>          <chr>    <dbl>     <dbl>    <dbl>     <dbl>
## 1 fixed    cond      <NA>           (Int…   0.0411   0.0247  -4.89e-3    0.0924
## 2 fixed    cond      <NA>           BMpe…   0.252    0.0415   1.73e-1    0.333
## 3 fixed    cond      <NA>           Refe…  -0.0689   0.0182  -1.04e-1   -0.0338
## 4 ran_pars cond      Location       sd__…   0.0333   0.0383   1.44e-3    0.138
## 5 ran_pars cond      Location:TankID sd__…   0.0108   0.00850  4.00e-4    0.0312
## 6 ran_pars cond      Residual       sd__…   0.128    0.00597  1.17e-1    0.140
```

This is the R2 of the model for dissolved fucoidan.

```
# R2 for fit3_dissolved
print(bayes_R2(fit3_dissolved))

##     Estimate  Est.Error      Q2.5     Q97.5
## R2 0.1742323 0.04065115 0.09675453 0.2548849
```

And here the model for SAF

```
# load model fit3_SAF or rerun
#fit3_SAF = brm(
#  bf(dCdt_sliding_mgLd_SAF ~ BMperV_gL + Reference_fucoidan_mgL_SAF +
#    (1 | Location/TankID)),
#  data = dat4saving[which(!dat4saving$Location %in% c("SA", "RO")),],
#  chains = 4, cores = 4, iter = 3000, warmup = 1000, thin = 2,
#  control = list(adapt_delta = 0.99))
#saveRDS(fit3_SAF, file = "20250620_model_fit3_SAF.rds")

fit3_SAF = readRDS("20250620_model_fit3_SAF.rds")

# Summary of the model fit3_SAF results
broom.mixed::tidy(fit3_SAF)

## # A tibble: 6 × 8
##   effect   component group          term  estimate std.error conf.low conf.high
##   <chr>    <chr>     <chr>          <chr>    <dbl>     <dbl>    <dbl>     <dbl>
## 1 fixed    cond      <NA>           (Int…  0.0122    0.0153  -1.71e-2    0.0419
## 2 fixed    cond      <NA>           BMpe…  0.0901    0.0167   5.76e-2    0.123
## 3 fixed    cond      <NA>           Refe… -0.0702    0.0188  -1.07e-1   -0.0341
## 4 ran_pars cond      Location       sd__…  0.0219    0.0234   1.44e-3    0.0849
## 5 ran_pars cond      Location:TankID sd__…  0.00513   0.00391  1.66e-4    0.0146
## 6 ran_pars cond      Residual       sd__…  0.0543    0.00272  4.93e-2    0.0599
```

with the corresponding R2

```
# R2 for fit3_SAF
print(bayes_R2(fit3_SAF))

##     Estimate  Est.Error      Q2.5     Q97.5
## R2 0.1785772 0.04170424 0.09876183 0.2612099
```

***Simulation of mesocosm experiment based on the model fit3***

To test the performance of the fit3 models predicting concentration changes over time based on brown algae biomass and standing fucoidan concentrations, this function simulates the conducted mesocosm experiment. The simulation uses epred_draws() to predict dC/dt from the posterior predictive distributions of the fit3 models fit3_dissolved and fit3_SAF. The simulated concentrations are plotted analogous to the observed values in Figure 2b and c (not shown here).

```
data_woSARO = dat4saving[which(!dat4saving$Location %in% c("SA", "RO")),]
SIM_tanks_slide = function(t, locations = 4, replicates = 3, dt = 2, Controls = FALSE){
  SIM_dat<-data.frame(Location = character(),
                      Replicate = numeric(),
                      Treatment = character(),
                      unique_tank = numeric(),
                      Day = numeric(),
                      Period = character(),
                      BMperV_gL = numeric(),
                      Reference_fucoidan_mgL_dissolved = numeric(),
                      Reference_fucoidan_mgL_SAF = numeric(),
                      dCdt_mgLdt_dissolved = numeric(),
                      dCdt_mgLdt_SAF = numeric(),
                      Fucoidan_mgL_dissolved = numeric(),
                      Fucoidan_mgL_SAF = numeric(),
                      Cumulative_fucoidan_mgL_dissolved = numeric(),
                      Cumulative_fucoidan_mgL_SAF = numeric())
  unique_tank = 0
  unique_BMV = 0
  for (loc in unique(data_woSARO$Location)){
    loc_tmp = randomNames(1, which.names = "first")
    for (rep in 1:replicates){
      if (Controls == TRUE){
        treatments = c("Algae", "Control")
      }
      else {
        treatments = "Algae"
      }
      for (treat_tmp in treatments){
        unique_tank = unique_tank + 1
        if (treat_tmp == "Algae"){
          unique_BMV = unique_BMV + 1
          BMV_tmp = unique(data_woSARO$BMperV_gL[data_woSARO$BMperV_gL > 0])[unique_BMV]
        }
        else {
          BMV_tmp = 0
        }
          for (time in seq(0,t,dt)){
            period_tmp = "synthesis"
            if (time == 0){
              ref_dissolved_tmp = 0
              ref_SAF_tmp = 0
            }
            else {
```

```
      ref_dissolved_tmp =
        as.numeric(SIM_dat$Fucoidan_mgL_dissolved[SIM_dat$Location == loc_tmp &
                                                  SIM_dat$Replicate == rep &
                                                  SIM_dat$Treatment == treat_tmp &
                                                  SIM_dat$Day == time-dt])
      ref_SAF_tmp =
        as.numeric(SIM_dat$Fucoidan_mgL_SAF[SIM_dat$Location == loc_tmp &
                                            SIM_dat$Replicate == rep &
                                            SIM_dat$Treatment == treat_tmp &
                                            SIM_dat$Day == time-dt])
    }
    if (time >= 13){
      BMV_tmp = 0
      period_tmp = "degradation"
    }
    SIM_dat_temp = data.frame(list(Location = loc_tmp, Replicate = rep,
                              Treatment = treat_tmp, unique_tank = unique_tank,
                              Day = time, Period = period_tmp,
                              BMperV_gL = BMV_tmp,
                              Reference_fucoidan_mgL_dissolved = ref_dissolved_tmp,
                              Reference_fucoidan_mgL_SAF = ref_SAF_tmp))
    colnames(SIM_dat_temp) = c("Location", "Replicate", "Treatment",
                              "unique_tank", "Day", "Period", "BMperV_gL",
                              "Reference_fucoidan_mgL_dissolved",
                              "Reference_fucoidan_mgL_SAF")
    SIM_dat_temp$Location = as.character(SIM_dat_temp$Location)
    SIM_dat_temp$Replicate = as.numeric(SIM_dat_temp$Replicate)
    SIM_dat_temp$Treatment = as.character(SIM_dat_temp$Treatment)
    SIM_dat_temp$unique_tank = as.numeric(SIM_dat_temp$unique_tank)
    SIM_dat_temp$Day = as.numeric(SIM_dat_temp$Day)
    SIM_dat_temp$Period = as.character(SIM_dat_temp$Period)
    SIM_dat_temp$BMperV_gL = as.numeric(SIM_dat_temp$BMperV_gL)
    SIM_dat_temp$Reference_fucoidan_mgL_dissolved =
      as.numeric(SIM_dat_temp$Reference_fucoidan_mgL_dissolved)
    SIM_dat_temp$Reference_fucoidan_mgL_SAF =
      as.numeric(SIM_dat_temp$Reference_fucoidan_mgL_SAF)
    dCdt_dissolved_tmp = epred_draws(fit3_dissolved, newdata = SIM_dat_temp,
                           ndraws = 1000, allow_new_levels = T) |>
      summarize(mean = mean(.epred))
    dCdt_SAF_tmp = epred_draws(fit3_SAF, newdata = SIM_dat_temp,
                           ndraws = 1000, allow_new_levels = T) |>
      summarize(mean = mean(.epred))
    if (time == 0){
      fucoidan_dissolved_tmp = rbeta(1, 2, 5)/2
      fucoidan_SAF_tmp = rbeta(1, 2, 5)/30
      cumulative_fucoidan_dissolved_tmp = fucoidan_dissolved_tmp
      cumulative_fucoidan_SAF_tmp = fucoidan_SAF_tmp
    }
    if (time > 0 & time <= 2){
      fucoidan_dissolved_tmp = ref_dissolved_tmp + dCdt_dissolved_tmp$mean*dt
      fucoidan_SAF_tmp = ref_SAF_tmp + dCdt_SAF_tmp$mean*dt
      cumulative_fucoidan_dissolved_tmp =
```

```
      SIM_dat$Cumulative_fucoidan_mgL_dissolved[SIM_dat$Location == loc_tmp &
                                              SIM_dat$Replicate == rep &
                                              SIM_dat$Treatment == treat_tmp &
                                              SIM_dat$Day == time-dt] +
      dCdt_dissolved_tmp$mean*dt
    cumulative_fucoidan_SAF_tmp =
      SIM_dat$Cumulative_fucoidan_mgL_SAF[SIM_dat$Location == loc_tmp &
                                        SIM_dat$Replicate == rep &
                                        SIM_dat$Treatment == treat_tmp &
                                        SIM_dat$Day == time-dt] +
      dCdt_SAF_tmp$mean*dt
  }
  if (time >2 & time <= 12){
    fucoidan_dissolved_tmp = ref_dissolved_tmp/2 + dCdt_dissolved_tmp$mean*dt
    fucoidan_SAF_tmp = ref_SAF_tmp/2 + dCdt_SAF_tmp$mean*dt
    cumulative_fucoidan_dissolved_tmp =
      SIM_dat$Cumulative_fucoidan_mgL_dissolved[SIM_dat$Location == loc_tmp &
                                              SIM_dat$Replicate == rep &
                                              SIM_dat$Treatment == treat_tmp &
                                              SIM_dat$Day == time-dt] +
      dCdt_dissolved_tmp$mean*dt
    cumulative_fucoidan_SAF_tmp =
      SIM_dat$Cumulative_fucoidan_mgL_SAF[SIM_dat$Location == loc_tmp &
                                        SIM_dat$Replicate == rep &
                                        SIM_dat$Treatment == treat_tmp &
                                        SIM_dat$Day == time-dt] +
      dCdt_SAF_tmp$mean*dt
  }
  if (time > 12){
    fucoidan_dissolved_tmp = ref_dissolved_tmp + dCdt_dissolved_tmp$mean*dt
    fucoidan_SAF_tmp = ref_SAF_tmp + dCdt_SAF_tmp$mean*dt
    cumulative_fucoidan_dissolved_tmp =
      SIM_dat$Cumulative_fucoidan_mgL_dissolved[SIM_dat$Location == loc_tmp &
                                              SIM_dat$Replicate == rep &
                                              SIM_dat$Treatment == treat_tmp &
                                              SIM_dat$Day == time-dt] +
      dCdt_dissolved_tmp$mean*dt
    cumulative_fucoidan_SAF_tmp =
      SIM_dat$Cumulative_fucoidan_mgL_SAF[SIM_dat$Location == loc_tmp &
                                        SIM_dat$Replicate == rep &
                                        SIM_dat$Treatment == treat_tmp &
                                        SIM_dat$Day == time-dt] +
      dCdt_SAF_tmp$mean*dt
  }
  SIM_dat = rbind(SIM_dat, c(loc_tmp, rep, treat_tmp, unique_tank,
                             time, period_tmp, BMV_tmp,
                             ref_dissolved_tmp, ref_SAF_tmp,
                             dCdt_dissolved_tmp$mean, dCdt_SAF_tmp$mean,
                             fucoidan_dissolved_tmp, fucoidan_SAF_tmp,
                             cumulative_fucoidan_dissolved_tmp,
                             cumulative_fucoidan_SAF_tmp))
  colnames(SIM_dat) = c("Location", "Replicate", "Treatment", "unique_tank",
```

```
                                 "Day", "Period", "BMperV_gL",
                                 "Reference_fucoidan_mgL_dissolved",
                                 "Reference_fucoidan_mgL_SAF",
                                 "dCdt_mgLdt_dissolved", "dCdt_mgLdt_SAF",
                                 "Fucoidan_mgL_dissolved", "Fucoidan_mgL_SAF",
                                 "Cumulative_fucoidan_mgL_dissolved",
                                 "Cumulative_fucoidan_mgL_SAF")
            SIM_dat$Location = as.character(SIM_dat$Location)
            SIM_dat$Replicate = as.numeric(SIM_dat$Replicate)
            SIM_dat$Treatment = as.character(SIM_dat$Treatment)
            SIM_dat$unique_tank = as.numeric(SIM_dat$unique_tank)
            SIM_dat$Day = as.numeric(SIM_dat$Day)
            SIM_dat$Period = as.character(SIM_dat$Period)
            SIM_dat$BMperV_gL = as.numeric(SIM_dat$BMperV_gL)
            SIM_dat$Reference_fucoidan_mgL_dissolved =
              as.numeric(SIM_dat$Reference_fucoidan_mgL_dissolved)
            SIM_dat$Reference_fucoidan_mgL_SAF =
              as.numeric(SIM_dat$Reference_fucoidan_mgL_SAF)
            SIM_dat$dCdt_mgLdt_dissolved = as.numeric(SIM_dat$dCdt_mgLdt_dissolved)
            SIM_dat$dCdt_mgLdt_SAF = as.numeric(SIM_dat$dCdt_mgLdt_SAF)
            SIM_dat$Fucoidan_mgL_dissolved = as.numeric(SIM_dat$Fucoidan_mgL_dissolved)
            SIM_dat$Fucoidan_mgL_SAF = as.numeric(SIM_dat$Fucoidan_mgL_SAF)
            SIM_dat$Cumulative_fucoidan_mgL_dissolved =
              as.numeric(SIM_dat$Cumulative_fucoidan_mgL_dissolved)
            SIM_dat$Cumulative_fucoidan_mgL_SAF =
              as.numeric(SIM_dat$Cumulative_fucoidan_mgL_SAF)
          }
        }
      }

  }
  return(SIM_dat)
}
sliding_sim = SIM_tanks_slide(24, locations = 4, replicates = 3, dt = 2, Controls = T)
```

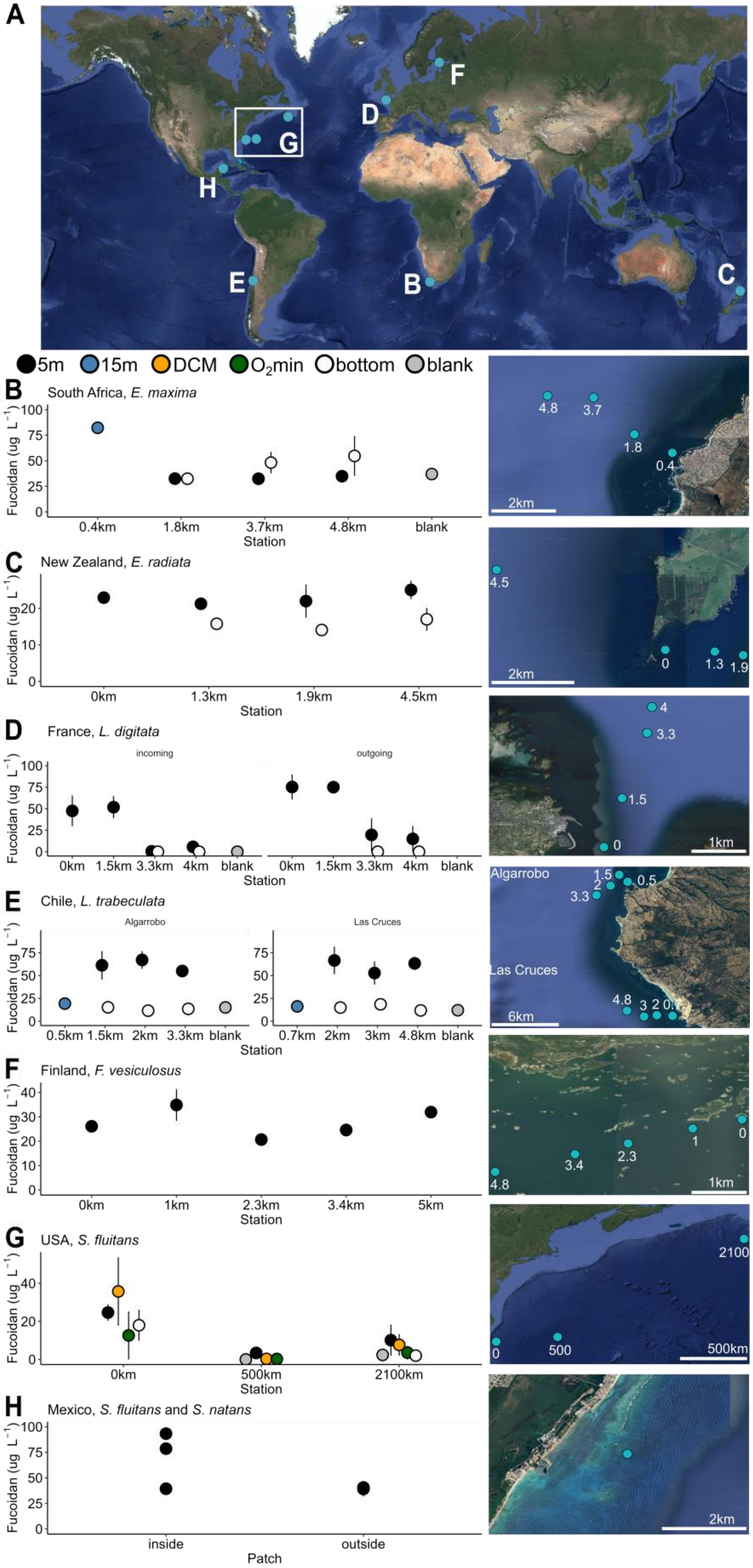
A
F
D
G
H
E
B
C
5m 15m DCM O2min bottom blank
B South Africa, E. maxima
Fucoidan (ug L−1)
0.4km 1.8km 3.7km 4.8km blank
Station
C New Zealand, E. radiata
0km 1.3km 1.9km 4.5km
D France, L. digitata
incoming
outgoing
0km 1.5km 3.3km 4km blank
E Chile, L. trabeculata
Algarrobo
Las Cruces
0.5km 1.5km 2km 3.3km blank
0.7km 2km 3km 4.8km blank
F Finland, F. vesiculosus
0km 1km 2.3km 3.4km 5km
G USA, S. fluitans
0km 500km 2100km
H Mexico, S. fluitans and S. natans
inside outside
Patch
2km
1km
6km
500km

**Fig. S1. Fucoidan concentrations (ug L$^{-1}$) after size and charge selection via anion exchange purification (AEX) in surface and bottom waters moving away from coastline.** Water samples taken (n=3 for each station) at depths of 5m (black), 15m (blue), deep chlorophyll maximum (yellow), $O_2$ minimum zone (green), bottom (white) and MilliQ blank samples processed with samples (grey) on the right plus sample maps on the left. Points represent the mean of three replicates with error bars indicating standard errors. **A)** Overview map of all sampling locations. **B)** South Africa, *E. maxima*, 0.4km away from coast sampled at 15m, 1.8km: 5m and 30m, 3.7km: 5m and 45m, 4.8km: 5m and 60m; **C)** New Zealand*, E. radiata* sampled at surface and bottom 0 to 4.5km away from algae bed; **D)** France, *L. digitata*, sampled during incoming and outgoing tides at four different stations in 5m and 30m depths (0 to 4km away from algae bed); **E)** Chile, *L. trabeculata*, Las Cruces, sampled at 0.5km away from coast at 15m, 1.5km away at 5m and 30m depth, 2 km away at 5m and 60m depth and 3.3 km away at 5m and 90m depth. In Algarrobo sampled at 15m 0.7km away from coast, at 5m depth and 30m depth 2km away, at 5m depth and 60m depth 3km away and at 5m depth and 90m depth 4.8km away from algae bed; **F)** Finland surface water sampling moving away up to 5 km from *Fucus vesiculosus* bed; **G)** USA, *S. fluitans*, on board R/V *Endeavor*, stations 21 -23, sampled 5m depth, DCM, $O_2$ min and bottom. Station 21 is directly off the continental shelf, around *S. fluitans* (0km), station 22 around 2100 km away and station 23 around 500km away from *S. fluitans*; **H)** Mexico, *S. fluitans* and *S. natans*, surface water samples taken inside and outside (around 50m away) of *Sargassum* patches. Fucoidan concentrations based on fucose concentrations of fucoidan standards extracted from biomass, except for *S. latissima* concentrations, which are calculated based on 30% of fucose.

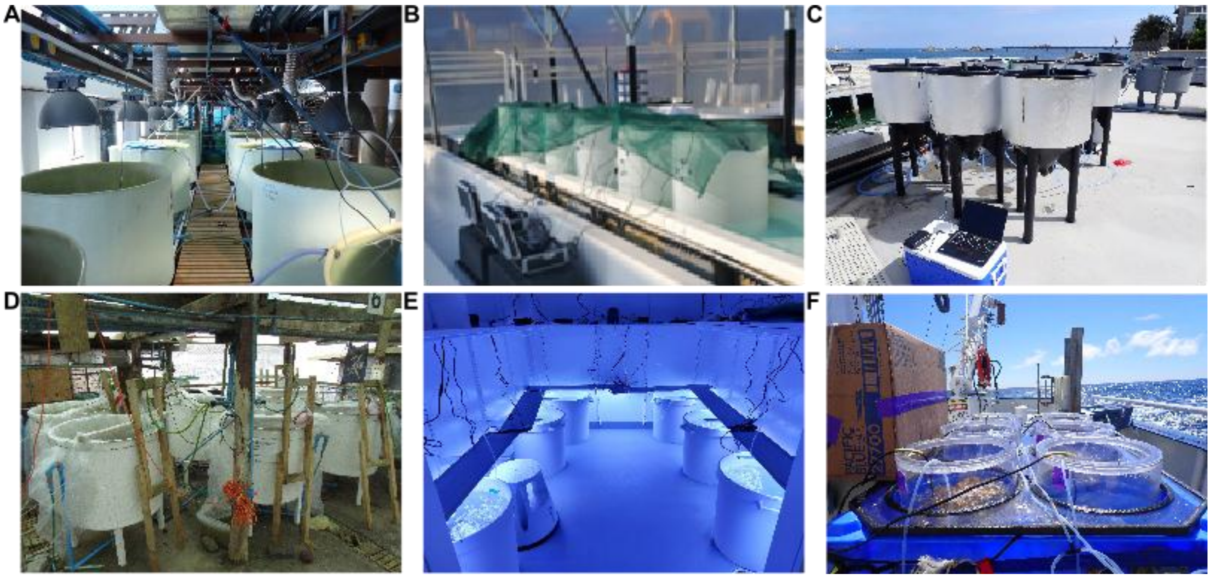


**Fig. S2. Tank systems of brown algae mesocosms. A)** *Ecklonia maxima* in South Africa, **B)** *Ecklonia radiata* in New Zealand, **C)** *Laminaria digitata* in France, **D)** *Lessonia trabeculata* in Chile, **E)** *Fucus vesiculosus* in Finland and **F)** *Sargassum fluitans* in the western North Atlantic, USA.

A

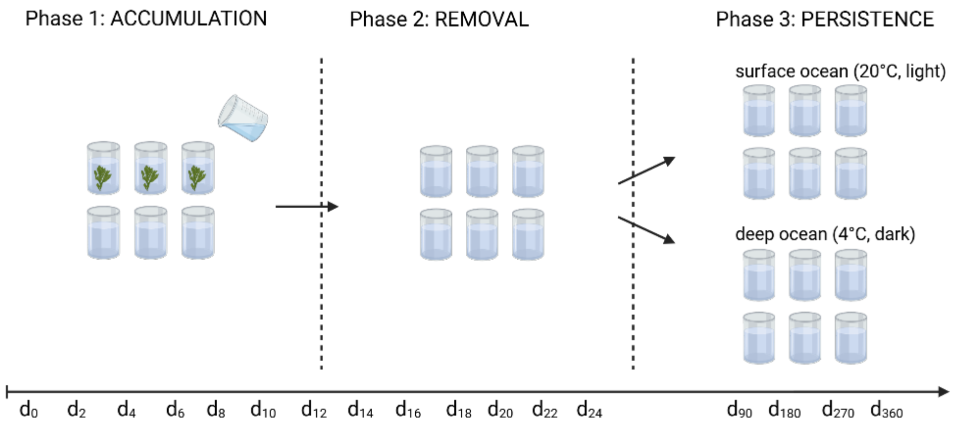

Phase 1: ACCUMULATION
Phase 2: REMOVAL
Phase 3: PERSISTENCE
surface ocean (20°C, light)
deep ocean (4°C, dark)
d0 d2 d4 d6 d8 d10 d12 d14 d16 d18 d20 d22 d24 d90 d180 d270 d360


B

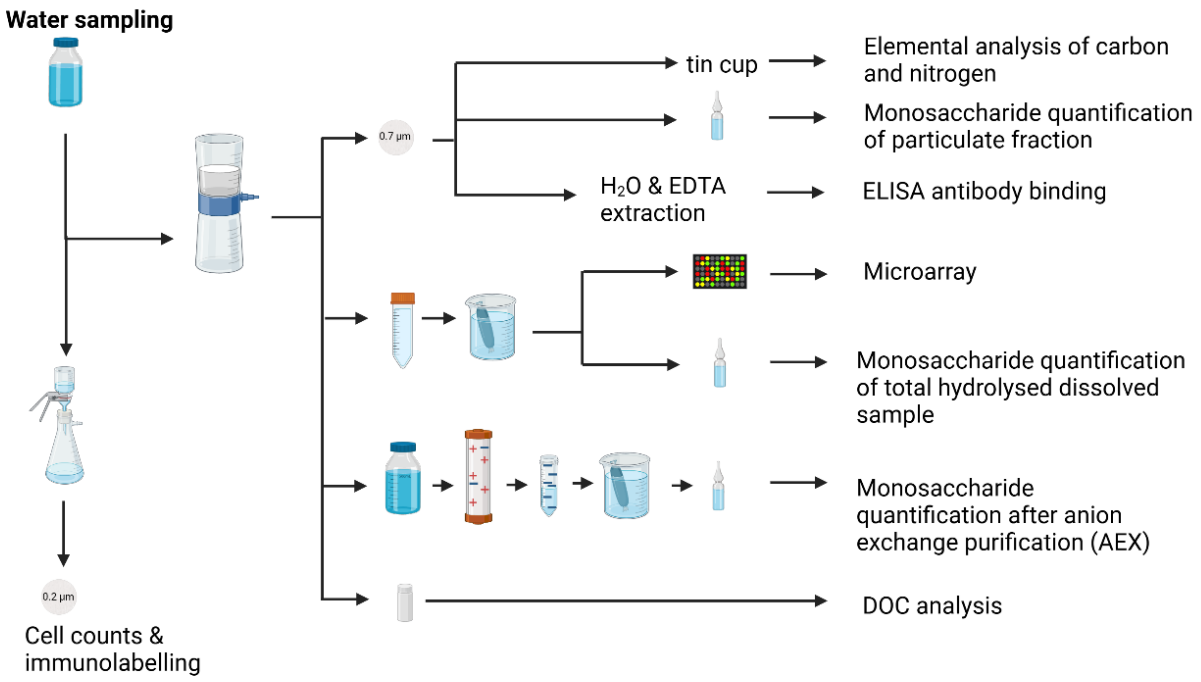

Water sampling
tin cup
Elemental analysis of carbon and nitrogen
0.7 µm
Monosaccharide quantification of particulate fraction
H2O & EDTA extraction
ELISA antibody binding
Microarray
Monosaccharide quantification of total hydrolysed dissolved sample
Monosaccharide quantification after anion exchange purification (AEX)
DOC analysis
0.2 µm
Cell counts & immunolabelling


C

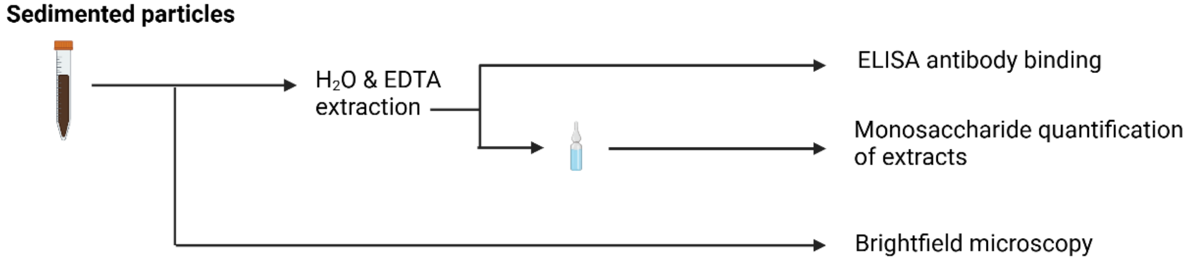

Sedimented particles
H2O & EDTA extraction
ELISA antibody binding
Monosaccharide quantification of extracts
Brightfield microscopy

**Fig. S3. Sampling scheme of brown algae incubations.** Created in BioRender [65]. **A)** Tank set-up consisting of six tanks. In phase 1 of experiment (Accumulation) three tanks were incubated with algae and three tanks served as controls for 12 days. Water was sampled every second day starting at day 0. After every water sampling half of the water was replenished. After 12 days algae were removed, starting phase 2 of experiment (Removal), in which water of algae and control tanks was kept for another 12 days. Water was sampled every second day. After 24 days, in phase 3 of experiment (Persistence) remaining water inside the tank was split up into two separate incubations for each tank. The first set was incubated at room temperature under a regulated light cycle, the second set was incubated at 4°C in the dark with the addition of nutrients. Samples were taken every 3 months for one year. **B)** At each sampling time 1 L of water was taken from each tank, and filtered through a pre-combusted 0.7 µm GFF filter. The filter was cut and prepared for elemental analysis, monosaccharide quantification of particulate fraction and specific antibody BAM1 ELISA binding. Filtered water was dialyzed and monosaccharides were quantified with HPAEC-PAD after acid hydrolysis. Polysaccharides were detected via antibody microarray binding. The surface active fucoidan (SAF) fraction was purified via anion exchange chromatography (AEX), dialyzed, acid hydrolyzed and monosaccharides quantified with HPAEC-PAD. DOC analysis was performed on acidified filtered water. Water was filtered through 0.2 µm polycarbonate filter for DAPI cell counts and BAM1 immunolabeling. **C)** Sedimented particles were scooped from bottom of the tanks after 24 days of incubation of *L. trabeculata*, *E. maxima* and *L. digitata* and sequentially extracted using MilliQ-water and 0.3M EDTA, monosaccharide quantification and specific antibody BAM1 ELISA binding was performed on extracts. Particles were imaged using a stereo microscope.

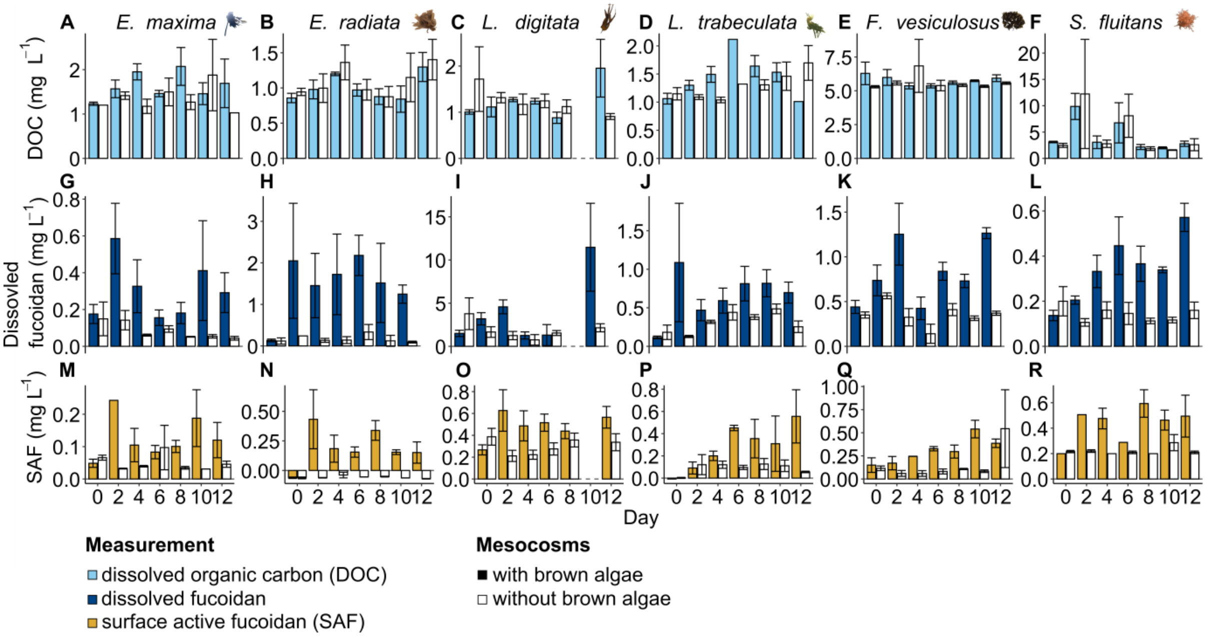


**Fig. S4. Fucoidan concentrations by all tested brown algae over 12 days of incubation. A-F)** Dissolved organic carbon (DOC) concentrations (mg $L^{-1}$), **G-L)** dissolved fucoidan concentrations (mg $L^{-1}$), **M-R)** and surface active fucoidan (SAF) concentrations (mg $L^{-1}$) during 12-day incubations in mesocosms with brown algae (filled) and without brown algae (not filled). Bars represent the mean of three mesocosm replicates with error bars indicating standard errors.

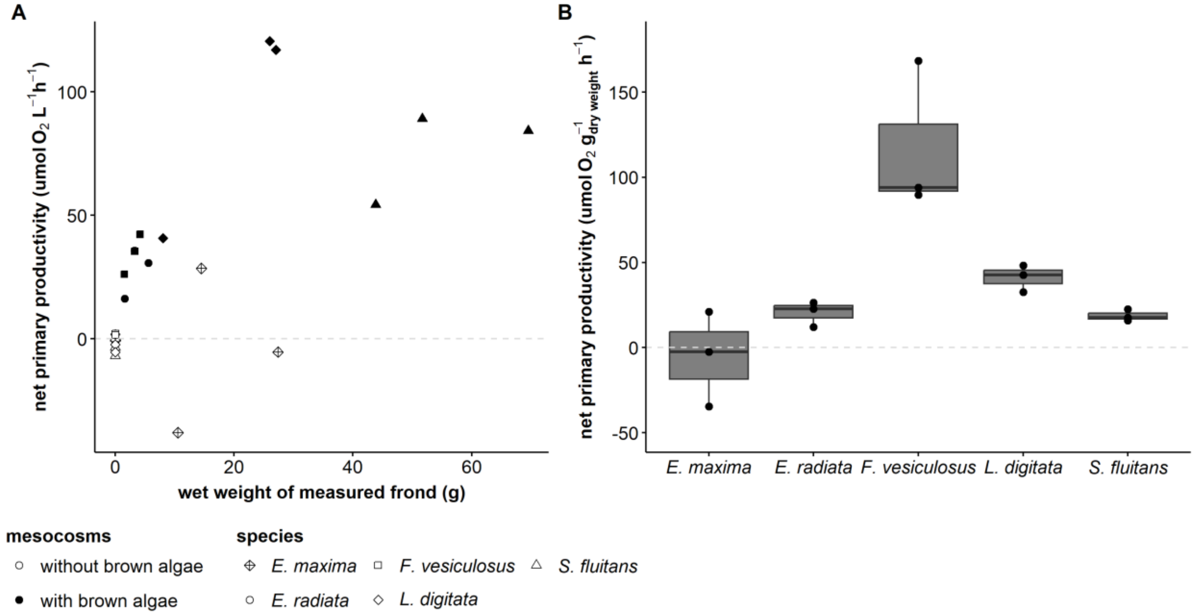


**Fig. S5. Net primary production calculated and scaled with incubated biomass. A)** Oxygen production in light and respiration in dark converted to net primary productivity and plotted against wet weight of the frond incubated to measure oxygen evolution. Data points at zero-intercept of x-axis mark mesocosms without brown algae. **B)** Net primary production per g dry weight of the incubated frond plotted by species. No data on oxygen production or consumption data are available for the incubation of *Lessonia trabeculata*.

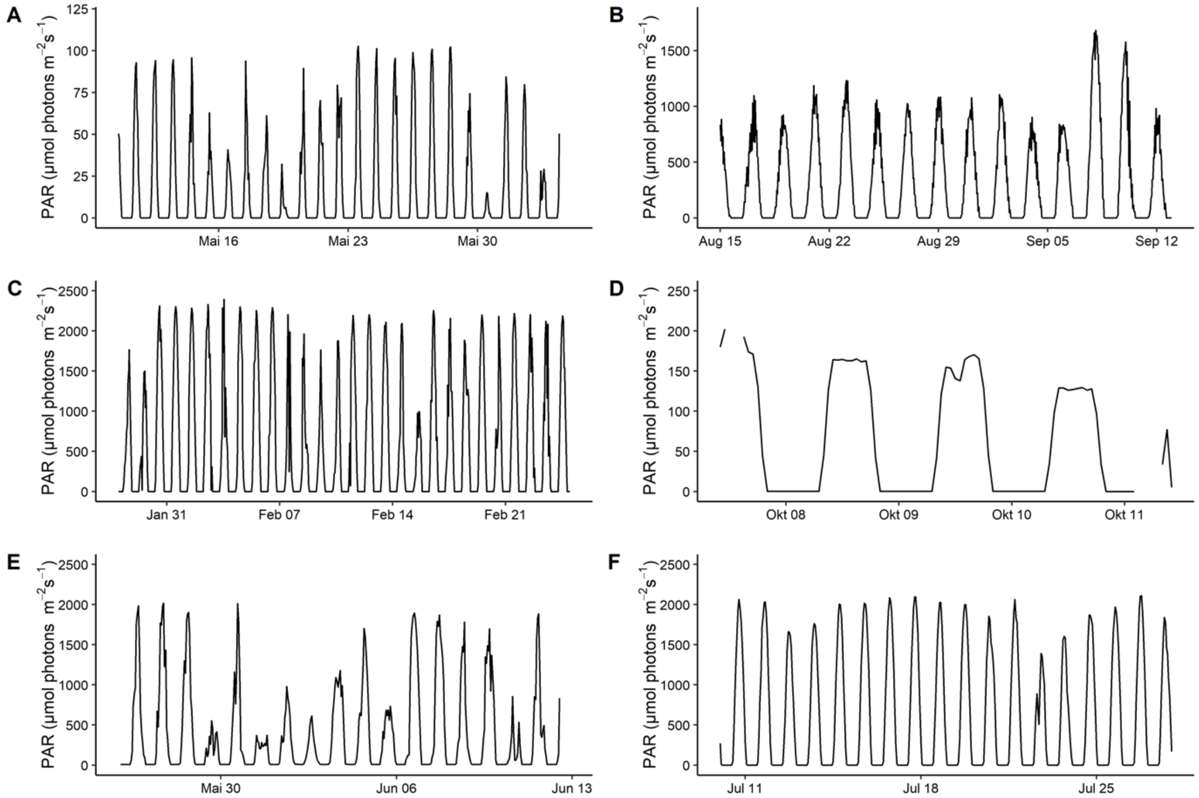


**Fig. S6. Hourly means of photosynthetically active radiation (PAR) during brown algae incubations. A)** PAR measurements were conducted using HOBO loggers in each tank during *Ecklonia radiata* incubations, New Zealand. **B)** Estimates for PAR during the *Laminaria digitata* incubation experiment at Roscoff, France, were calculated from NASA POWER (https://power.larc.nasa.gov/) all sky surface total PAR. **C)** Radiation data for the *Lessonia trabeculata* incubation from Las Cruces, taken from the meteorological station of the Centro de Estudios Avanzados en Zonas Áridas (CEAZA), Chile. **D)** A RBRsolo logger with an attached PAR-sensor (LI-192 Underwater Quantum Sensor) was used to measure PAR in one tank during *Fucus vesiculosus* incubations. **E)** The PAR data during the *Sargassum fluitans* incubation was recorded by R/V *Endeavor*. **F)** Estimates for PAR during *Sargassum fluitans* bottle incubations at Puerto Morelos, Mexico, were calculated from NASA POWER (https://power.larc.nasa.gov/) all sky surface total PAR. No PAR data is available for *S. latissima* algae farm sampling and for *Ecklonia maxima* mesocosms experiment irradiated with an Osram Power Star hqi-e 440W/D lamp located centrally above mesocosms with a nominal flux of 40000 lumen for 14 hours daily.

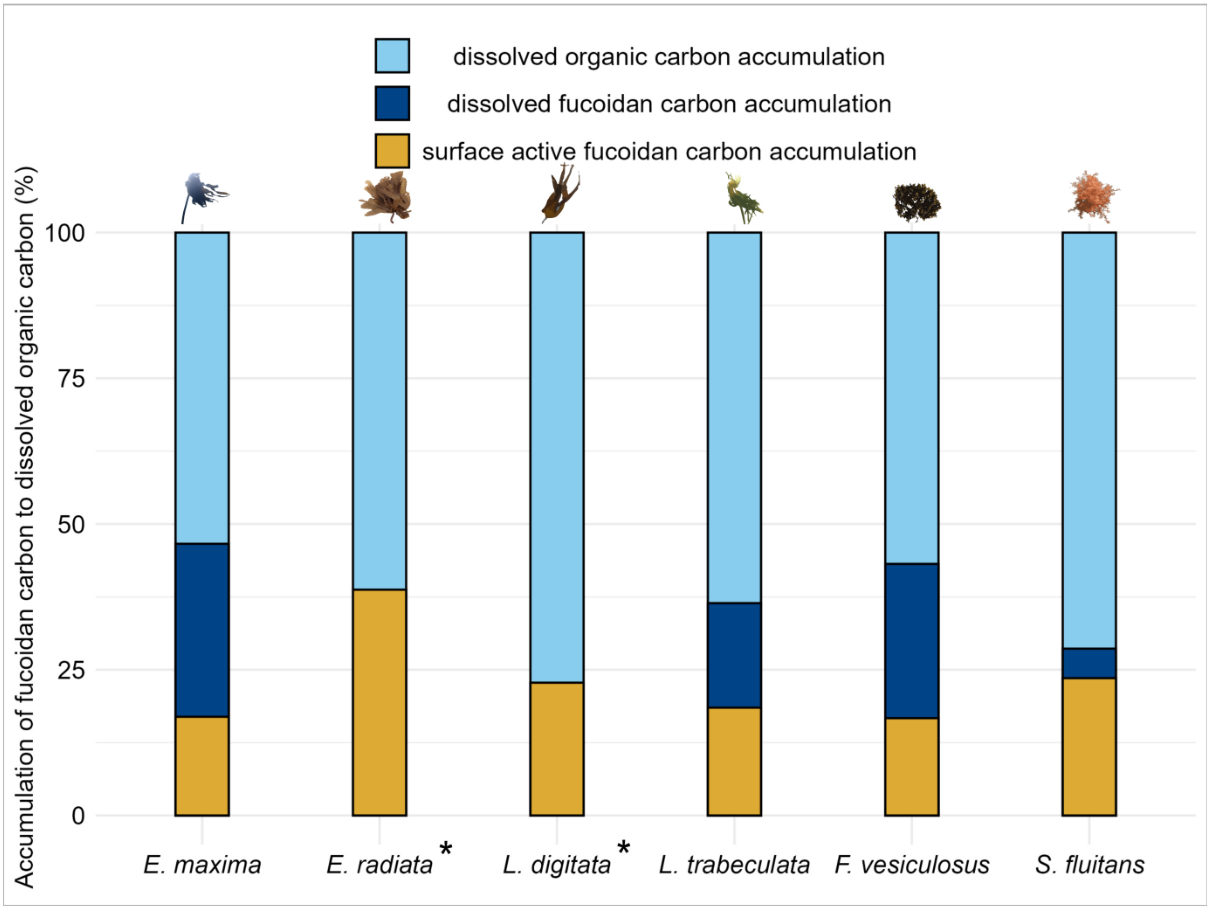


**Fig. S7. Up to 50% of total dissolved organic carbon (DOC) can be attributed to dissolved fucoidan carbon. Up to 30% confirmed by surface active fucoidan (SAF) fraction.** DOC (light blue) concentrations were set to 100% to calculate relative fucoidan carbon accumulation. Both fucoidan fractions refer to the quantified fucose in bulk hydrolysable polymers > 1kDa without (dark blue) and with specific anion exchange chromatography, targeting the surface active fucoidan fraction (yellow). * indicates those species not fully-assessed due to excluded dissolved fucoidan estimates that were greater than quantified DOC. Fucoidan carbon comprised the following proportions of DOC: *E. maxima*: 47% dissolved fucoidan, 17% SAF; *L. digitata*: 23% SAF; *E. radiata*: 39% SAF; *L. trabeculata:* 36% dissolved fucoidan, 18% SAF; *F. vesiculosus*: 43% dissolved fucoidan, 16% SAF; *S. fluitans:* 29% dissolved fucoidan, 24% SAF (DOC concentrations above 500 µM were excluded for *S. fluitans* incubations due to contamination).

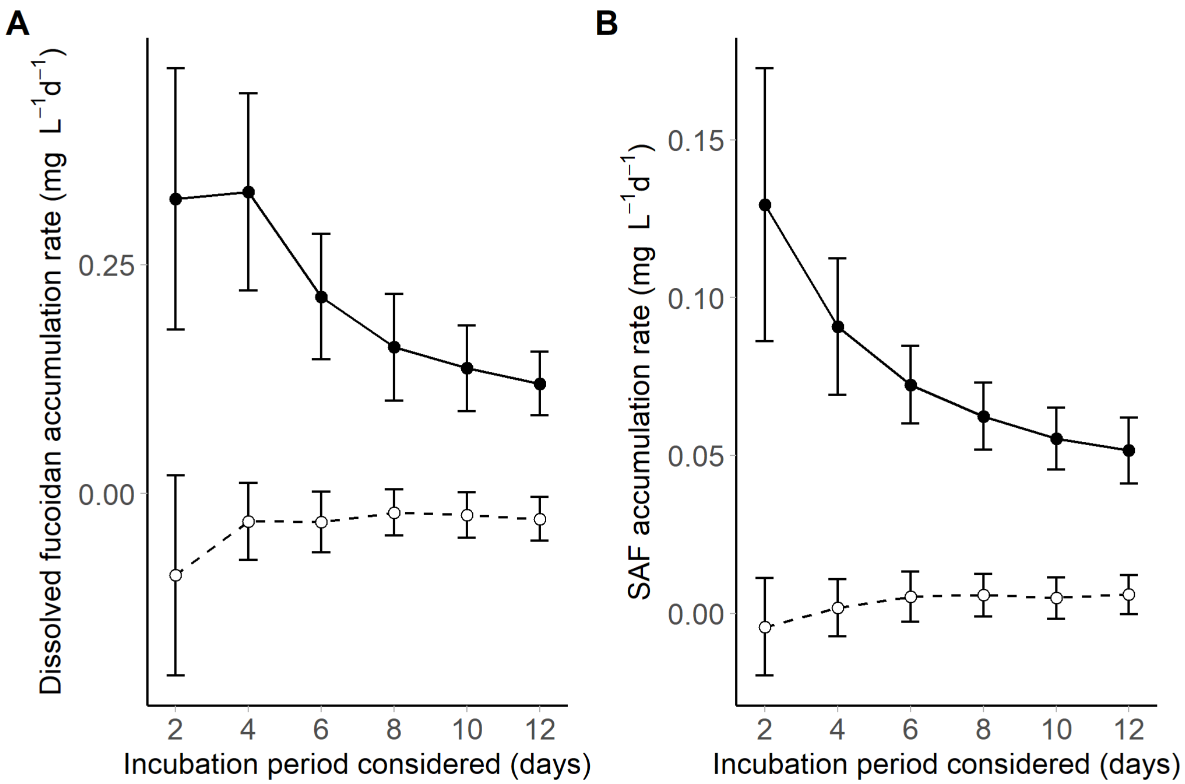


**Fig. S8. Fucoidan accumulation rates peak during the first four days of incubation.** Fucoidan accumulation rate ± s.e.m. (n=18) in mesocosms with brown algae (black, solid line) and mesocosms without brown algae (white, dashed line) in mg L$^{-1}$ d$^{-1}$ over 12 days of incubation for **A)** dissolved fucoidan and **B)** surface active fucoidan (SAF).

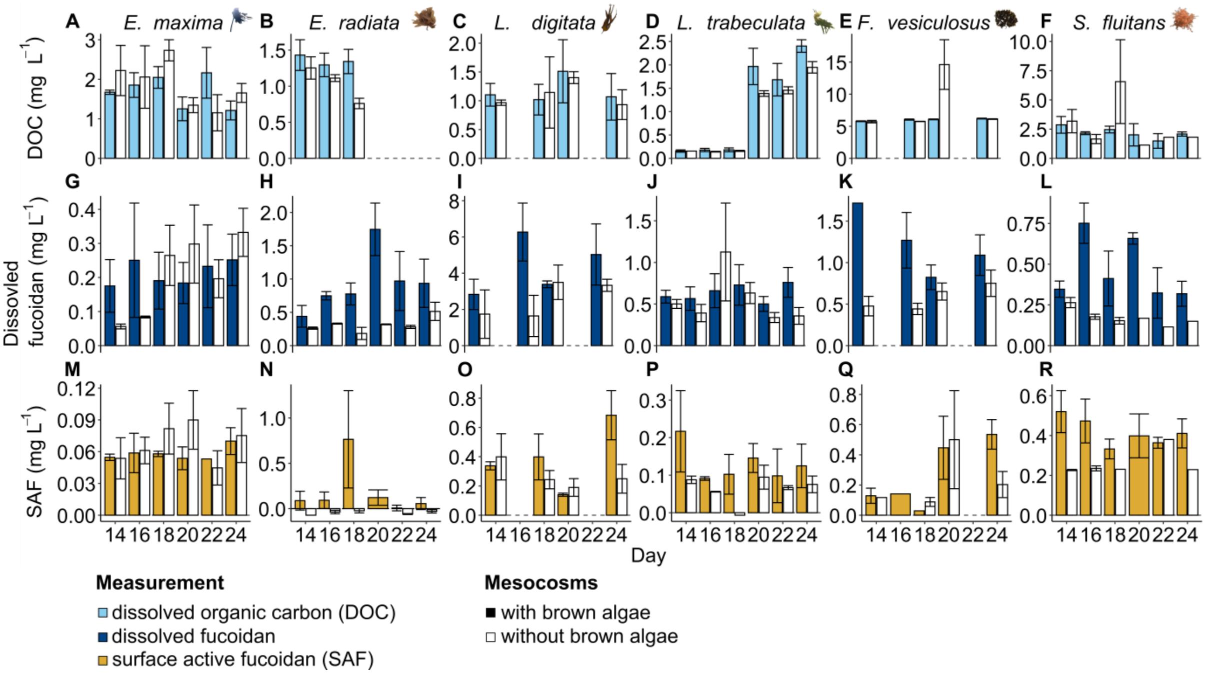


**Fig. S9. Fucoidan concentrations by all tested brown algae over 12 days of incubation after removal of brown algae (day 14 to 24). A-F)** Dissolved organic carbon (DOC) concentrations (mg $L^{-1}$), **G-L)** dissolved fucoidan concentrations (mg $L^{-1}$), **M-R)** and surface active fucoidan (SAF) concentrations (mg $L^{-1}$) during 12-day incubations after removal of brown algae in mesocosms with brown algae (filled) and without brown algae (not filled). Bars represent the mean of three mesocosm replicates with error bars indicating standard errors.

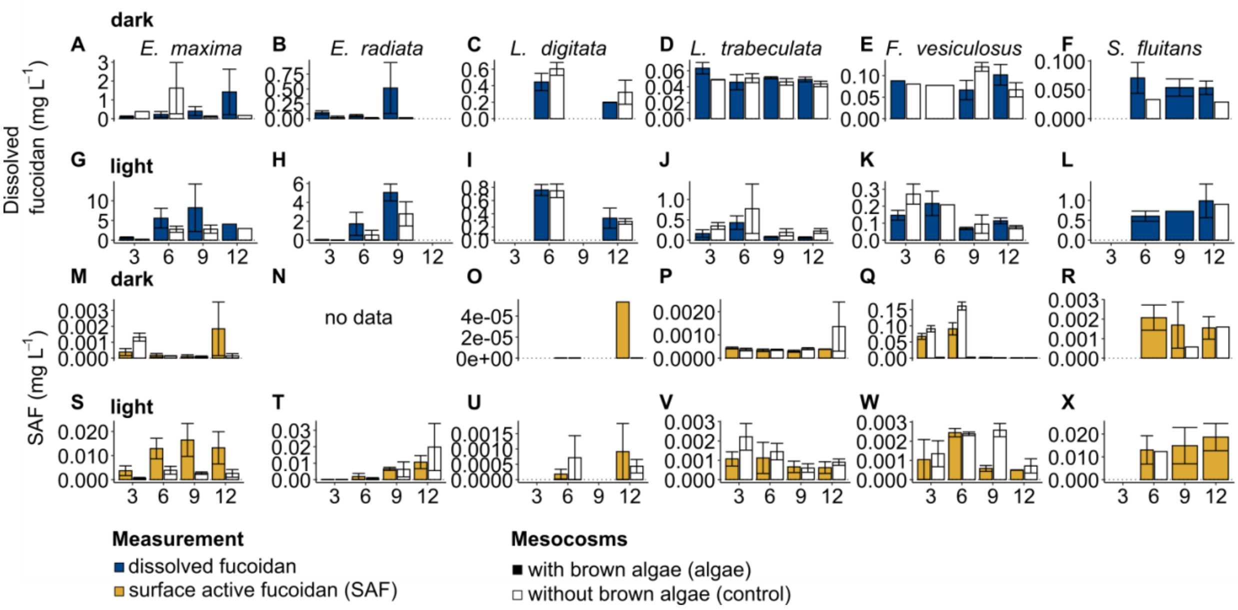


**Fig. S10. Fucoidan concentrations (mg L$^{-1}$) by all tested brown algae over year long incubation (3-12 months). A-F)** Dissolved fucoidan during dark incubations and **G-L)** light incubations and **M-R)** surface active fucoidan (SAF) concentrations during dark and **S-X)** light incubations in mesocosms with brown algae (filled bars) and without brown algae (not filled bars). Bars represent the mean of three mesocosm replicates with error bars indicating standard errors. **N)** No data available for *E. radiata* incubation, quantified based on SAF fraction.

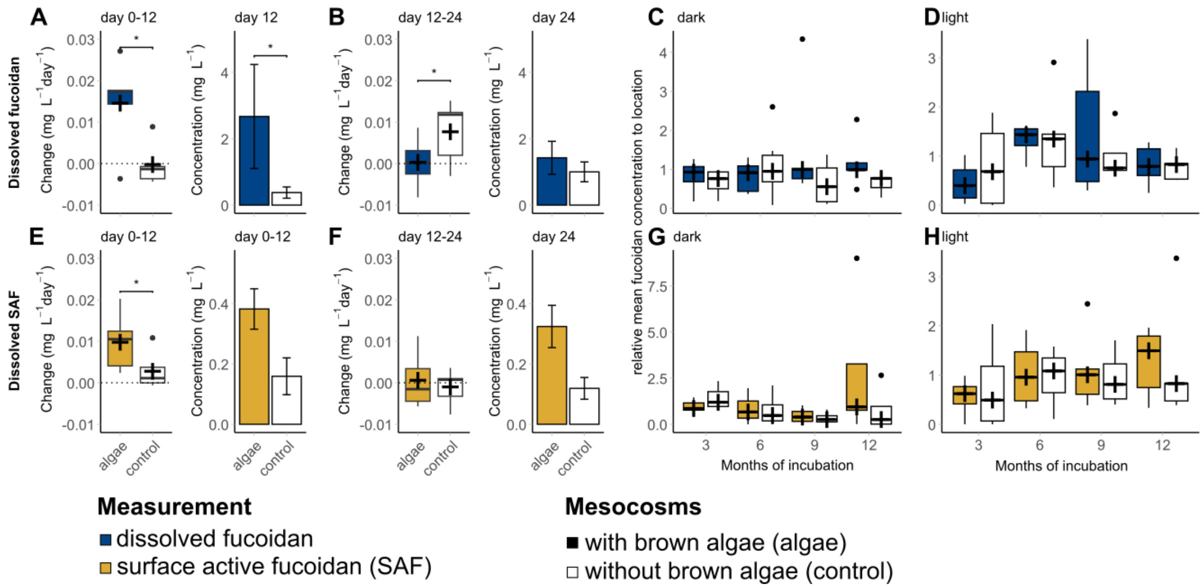


**Fig. S11. Detectable fucose polymers after a year-long incubation**. For **dissolved fucoidan estimates (A-D)**, **A)** fucoidan concentrations (mg $L^{-1}$ $d^{-1}$) increased significantly between algae and controls from day 0 to day 12 ($p = 0.03125$, Wilcoxon-Test). On day 12, algae had higher fucoidan concentrations (± s.e.m, mg $L^{-1}$) than controls ($p = 0.03125$ vs. $p = 0.07278$, Wilcoxon-Test). **B)** After algae removal (day 12 to day 24), fucoidan concentrations increased significantly in control mesocosms ($p = 0.03125$, Wilcoxon-Test), with no significant differences between algae and control tanks on day 24 ($p = 0.0625$, Wilcoxon-Test). Long-term analysis of **C)** dark- and **D)** light-incubated mesocosm water sampled every 3 months for a year showed no significant differences (Wilcoxon-Test). For **dissolved surface active fucoidan (SAF) (E-H)**, **E)** a significant increase in fucoidan concentrations occurred between algae and controls from day 0 to day 12 ($p = 0.03125$, Wilcoxon-Test), but no significant differences were detected on day 12 ($p = 0.07278$, Wilcoxon-Test). **F)** Between days 12 and 24, no significant changes were observed in fucoidan concentrations ($p = 0.8438$, Wilcoxon-Test), and no significant differences were found on day 24 ($p = 0.05072$, Wilcoxon-Test). Similarly, no significant differences were detected in **G)** dark- or **H)** light-incubated mesocosm water sampled every 3 months for one year (Wilcoxon-Test).

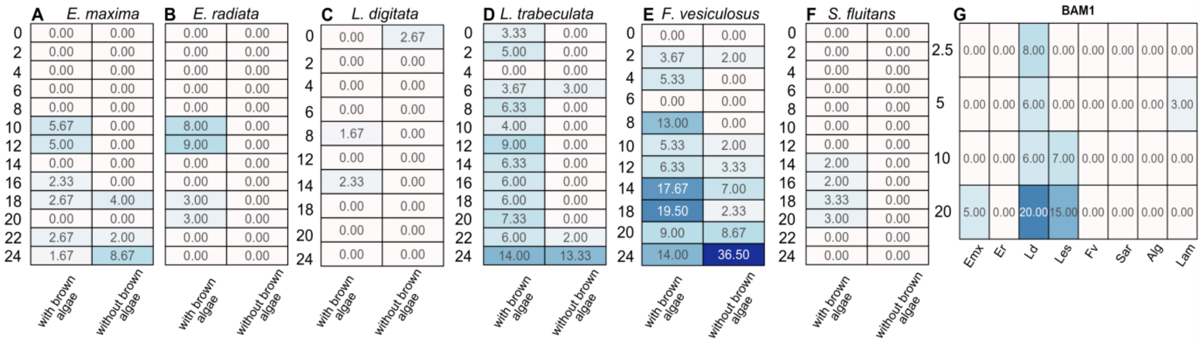


**Fig. S12. Fucoidan-specific monoclonal antibody BAM1 confirmed persistence of dissolved fucoidan in *L. trabeculata* mesocosms. A-F, G)** Signals of fucoidan-specific BAM1 in triplicate mesocosms with and without brown algae over 24 days. Each signal represents mean values of **A)** *E. maxima*, **B)** *E. radiata,* **C)** *L.digitata,* **D)** *L. trabeculate***, E)** *F. vesiculosus***, F)** *S. fluitans*, **G)** BAM1 binding to standard series of fucoidans (concentrations in mg $L^{-1}$ on left) extracted from biomass (Emx: *E. maxima* fucoidan, Er: *E. radiata* fucoidan, L.d: *L. digitata* fucoidan, Les: *L. trabeculata* fucoidan, F.v.: *F. vesiculosus* fucoidan*,* Sar: *S. fluitans* fucoidan) and alginate (ALG) and laminarin (LAM) indicate antibody specificity and detection ranges. Boxes are colored according to antibody signals.

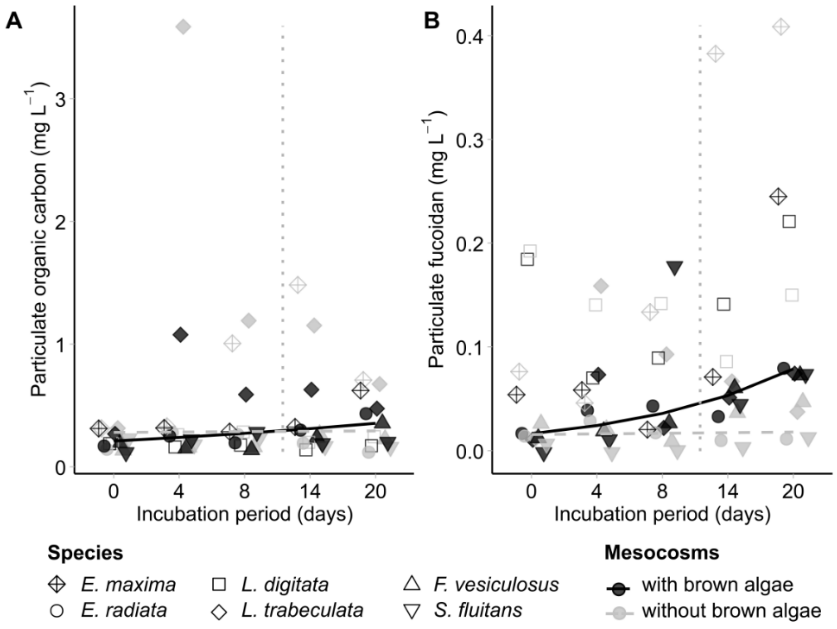


**Fig. S13. Increase of particulate organic carbon and particulate fucoidan from day 0 to day 20**. Stronger increase of **A)** particulate organic carbon (with brown algae: $R^2=0.064$, $p=0.147$; without: $R^2=-0.055$, $p=0.943$) and **B)** particulate fucoidan (with brown algae: $R^2=0.463$, $p< 0.001$; without: $R^2=0.062$, $p=0.944$) in mg $L^{-1}$ in mesocosms with brown algae (black) compared to mesocosms without brown algae (grey). Each data point represents the mean of 3 mesocosms. Calculations exclude *E. maxima* and *L. digitata.* The vertical dotted grey line indicates when algal biomass was removed.

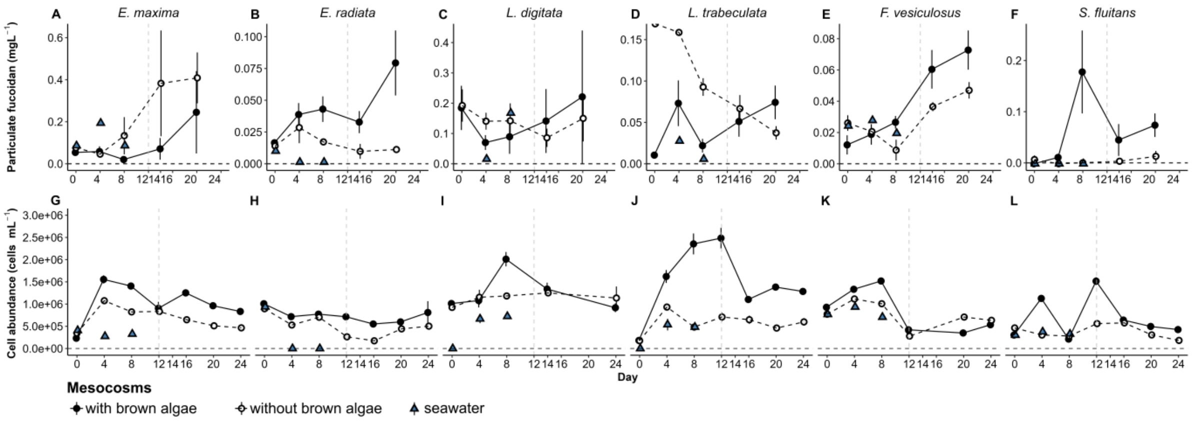


**Fig. S14. Fucoidan precipitation may affect microbial cell abundance. A-F)** Particulate fucoidan concentrations ± s.e.m. during mesocosms with brown algae (black circle, n=3) and mesocosms without brown algae (white circle, n=3) and fresh seawater (blue triangle, n=1). **G-L)** Microbial cell abundance estimated from DAPI-stained 0.2 µm polycarbonate filters for brown algae (black circle) and without brown algae (white circle), as well as fresh seawater (blue triangle). The vertical dotted grey line indicates when algal biomass was removed.

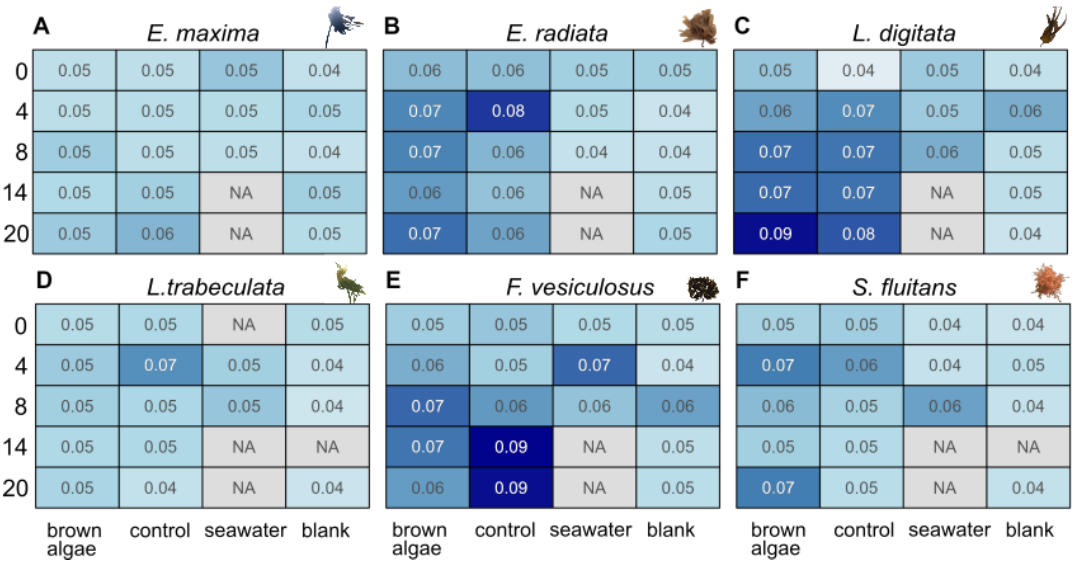


**Fig. S15. Fucoidan specific BAM1 antibody signal in particulate fraction in brown algae and without brown algae (control) incubations.** BAM1 signal in particulate fraction increases from day 0 to day 4 of all incubations, and is stable after day 4, measured for day 8, 14 and 20. Mesocosms with and without brown algae each represent n=3, seawater and blank each represent n=1. No difference in signal between brown algae and control tanks of the following incubations **A)** *E. maxima*, **B)** *E. radiata*, **C)** *L. digitata*, **D)** *L. trabeculata*, **E)** *F. vesiculosus* and **F)** *S. fluitans*.

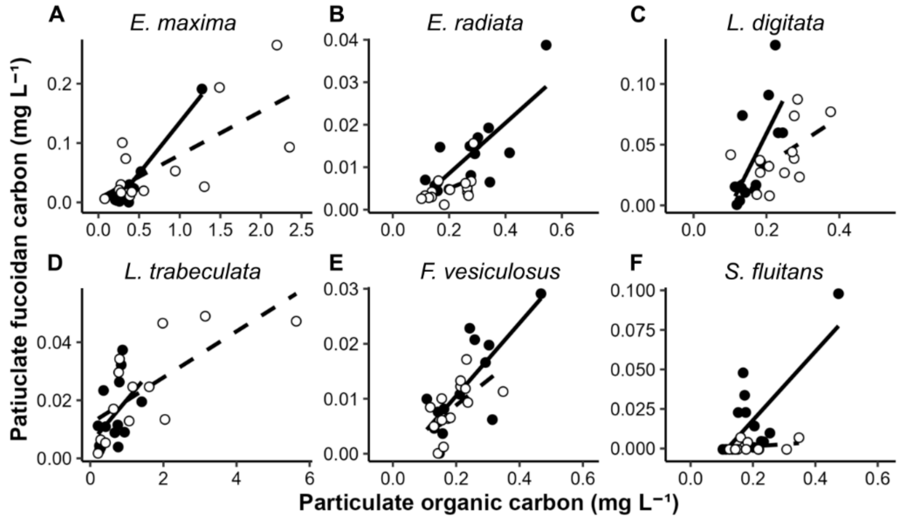


**Fig. S16. Spearman's correlation between particulate organic carbon and particulate fucoidan carbon with lines fitted with least-squares regression line.** Mesocosms with brown algae shown in black, solid line and mesocosms without brown algae in white, dashed line. **A**) *E. maxima* incubation (with brown algae: $R^2$=0.93, p<0.001, without brown algae: $R^2$= 0.48, p=0.003), **B)** *E. radiata* incubations (with brown algae: $R^2$=0.59, p<0.001; without brown algae: $R^2$=0.23, p=0.05), **C)** *L. digitata* incubations (with brown algae: $R^2$=0.42, p=0.009; without brown algae: $R^2$=0.24, p=0.05), **D)** *L. trabeculata* incubations (with brown algae: $R^2$=0.15, p=0.09; without brown algae: $R^2$=0.47, p=0.005), **E)** *F. vesiculosus* incubations (with brown algae: $R^2$=0.57, p<0.001; without brown algae: $R^2$=0.29, p=0.02), **F)** *S. fluitans* incubations (with brown algae: $R^2$=0.49, p=0.003; without brown algae: $R^2$=-0.001, p=0.35).

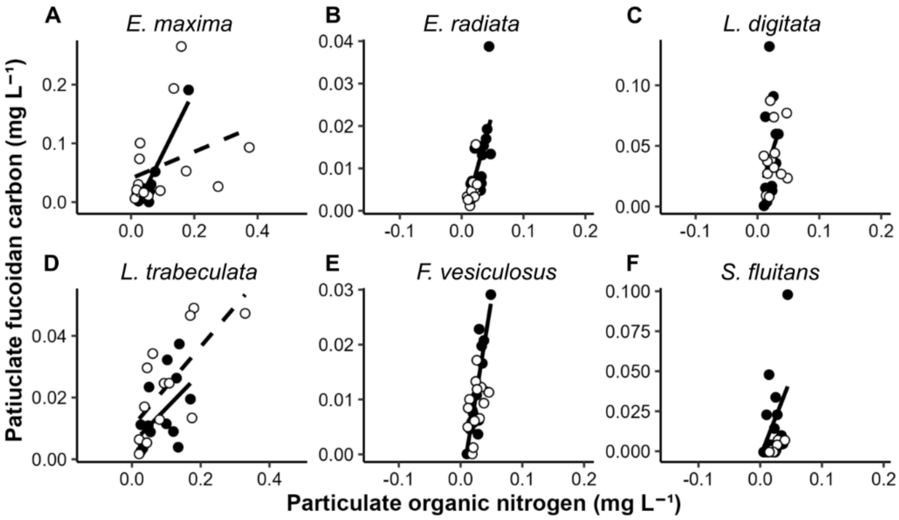


**Fig. S17. Spearman's correlation between particulate organic nitrogen and particulate fucoidan with lines fitted with least-squares regression line.** Mesocosms with brown algae shown in black, solid line and mesocosms without brown algae in white, dashed line. **A**) *E. maxima* incubation (with brown algae: $R^2$=0.87, p<0.001, without brown algae: $R^2$= 0.04, p=0.24), **B)** *E. radiata* incubations (with brown algae: $R^2$=0.33, p=0.02; without brown algae: $R^2$=0.24, p=0.04), **C)** *L. digitata* incubations (with brown algae: $R^2$=0.01, p=0.33; without brown algae: $R^2$=-0.04, p=0.49), **D)** *L. trabeculata* incubations (with brown algae: $R^2$=0.19, p=0.06; without brown algae: $R^2$=0.45, p=0.01), **E)** *F. vesiculosus* incubations (with brown algae: $R^2$=0.43, p<0.001; without brown algae: $R^2$=0.08, p=0.15), **F)** *S. fluitans* incubations (with brown algae: $R^2$=0.14, p=0.11; without brown algae: $R^2$=0.60, p=0.003).

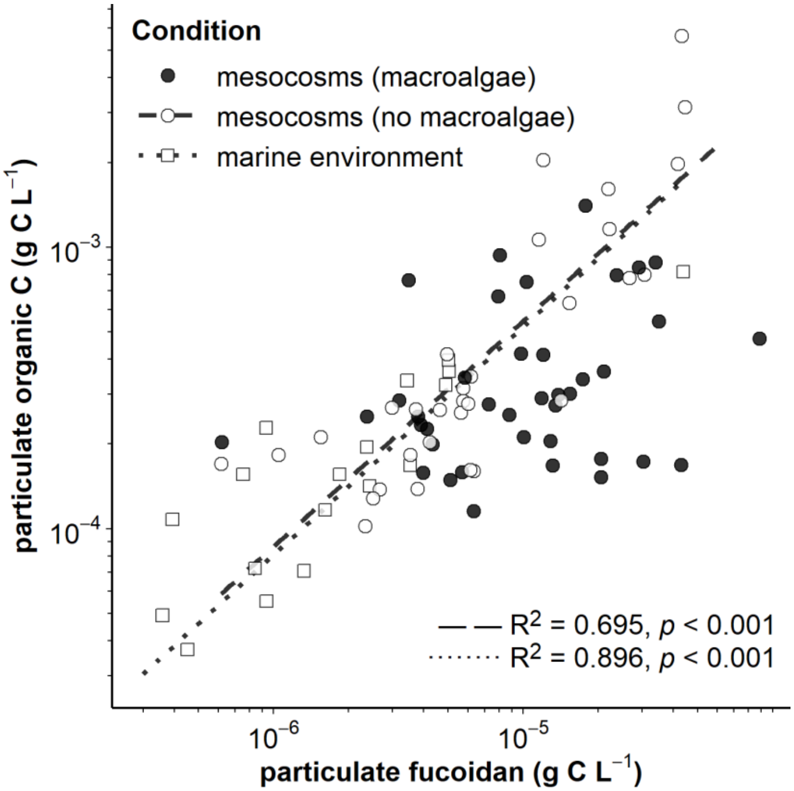


**Fig. S18. Power law relationship of fucoidan carbon from particulate fucose against particulate organic carbon (POC)** in brown algae mesocosms (black circles), mesocosms without brown algae (dashed line, white circles) and values inferred from previous studies without dedicated fucoidan quantification (dotted line, white squares).

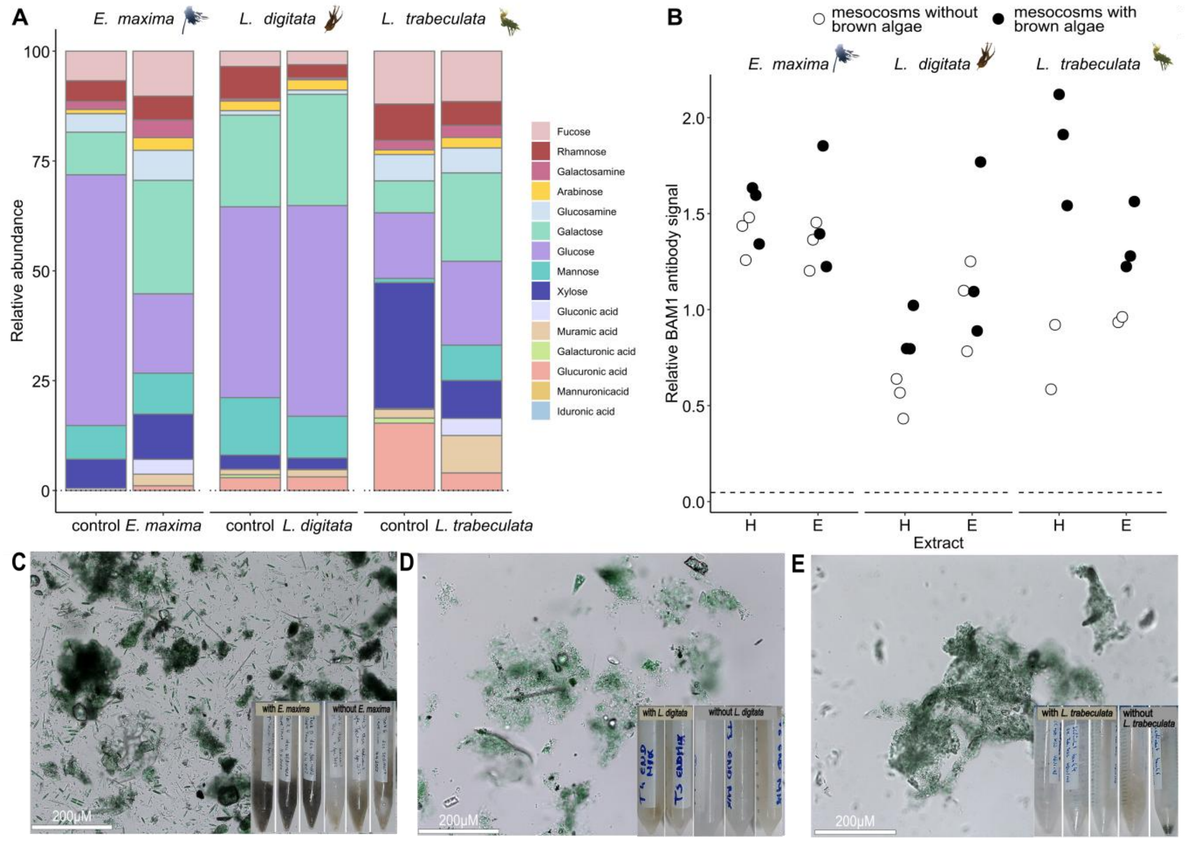


**Fig. S19. Fucoidan presence confirmed in sedimented particles.** Sedimented particles of *Ecklonia maxima, Laminaria digitata, Lessonia trabeculata* and control (without brown algae) mesocosms sampled on day 24 of incubations. **A)** Relative abundance of monosaccharide of particles, dominated by glucose and galactose. **B)** BAM1 specific antibody binding to water (H) and EDTA (E) extracts of particles in brown algae tanks, as well as tanks without brown algae showing presence of fucoidan in sedimented particles originating from macro- and microalgae. Brightfield image of sedimented particles and particulate sampled in **C)** *E. maxima*, **D)** *L. digitata* and **E)** *L. trabeculata* incubations**.**

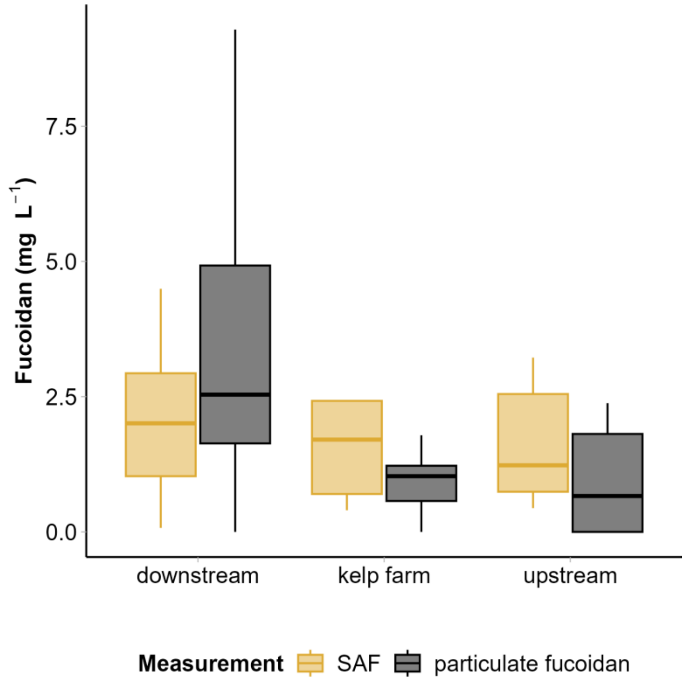


**Fig. S20. Fucoidan presence around kelp farm.** Surface active fucoidan (SAF; yellow) and particulate fucoidan (grey) in mg $L^{-1}$ upstream (n = 6), inside (n = 9) and downstream (n = 12) of *S. latissima* kelp farm.

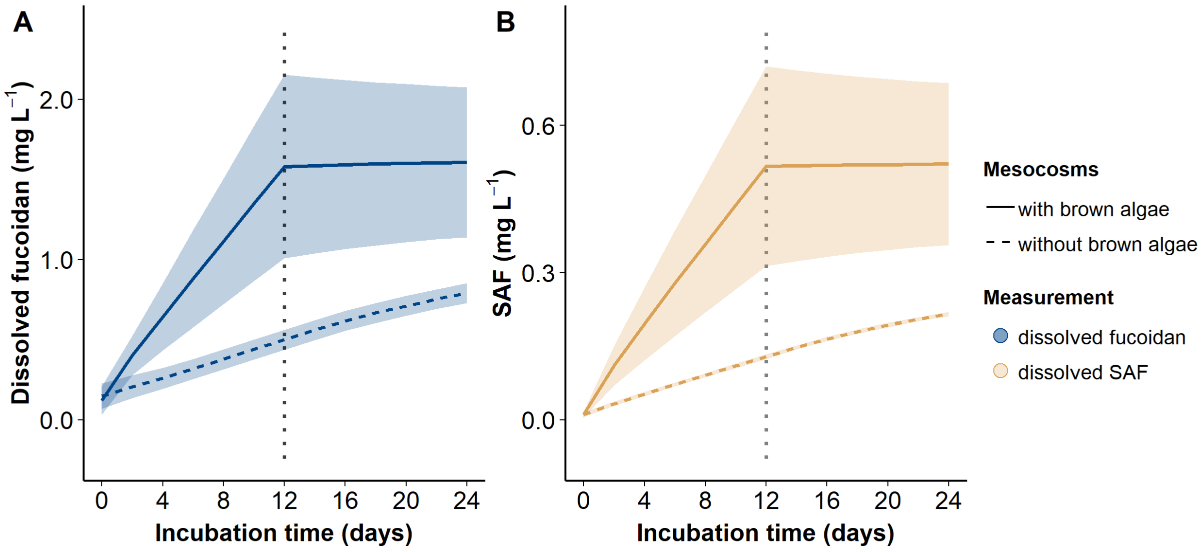


**Fig. S21. Posterior distributions of hierarchical Bayesian model recreate experimental observations in simulations of mesocosm experiments** for **A)** dissolved fucoidan and **B)** surface active fucoidan (SAF) in mg $L^{-1}$ for mesocosms with brown algae (solid line) and mesocosms without brown algae (dashed line) over 24 days of incubation. The dotted grey line at 12 days incubation time marks the removal of brown macroalgae from the mesocosms.

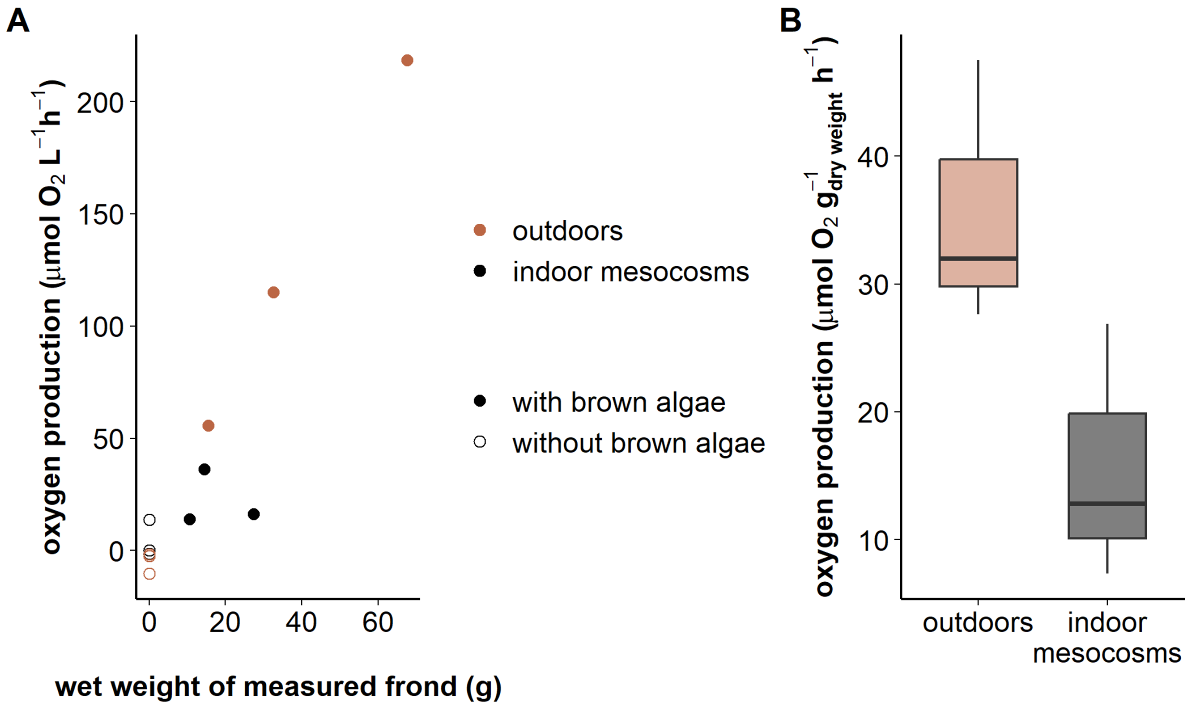


**Fig. S22. Substantially higher oxygen production of *in situ E. maxima* specimens compared to specimens kept indoor in tanks. A)** Oxygen production scaled with wet weight of incubated fronds for outdoor algae (red circles) but not for indoor algae (black circles) with incubations without brown algae in white. **B)** Oxygen production for outdoor algae in natural light exceeded oxygen production for tank algae under artificial light.

**Table S1.** Overview of environmental sampling, including species, country, location, station, estimated distance to algae, latitude and longitude.

| Species | Country | Location | Station | Distance to algae | Longitude | Latitude |
|---|---|---|---|---|---|---|
| *Ecklonia maxima* | South Africa | Seapoint | Stn1 | 0.4km | 33°55.200’ S | 18°22.405’ E |
| *Ecklonia maxima* | South Africa | Seapoint | Stn2 | 1.8km | 33°54.830’ S | 18°21.613’ E |
| *Ecklonia maxima* | South Africa | Seapoint | Stn3 | 3.7km | 33°54.033’ S | 18°20.783’ E |
| *Ecklonia maxima* | South Africa | Seapoint | Stn4 | 4.8km | 33°54.025’ S | 18°19.889’ E |
| *Ecklonia radidata* | New Zealand | Tauranga | Stn1 | 4.5km | 37°37'40.73"S | 176°21'46.84"E |
| *Ecklonia radidata* | New Zealand | Tauranga | Stn2 | 0km | 37°39'2.29"S | 176°24'35.08"E |
| *Ecklonia radidata* | New Zealand | Tauranga | Stn3 | 1.3km | 37°39'3.97"S | 176°25'24.94"E |
| *Ecklonia radidata* | New Zealand | Tauranga | Stn4 | 1.9km | 37°39'8.80"S | 176°25'53.90"E |
| *Laminaria digitata* | France | Roscoff | Stn1 - incoming | 0km | 48°42.878’ N | 3°57.355’W |
| *Laminaria digitata* | France | Roscoff | Stn2 - incoming | 1.5km | 48°43.637’ N | 3°57.091 W |
| *Laminaria digitata* | France | Roscoff | Stn3 - incoming | 3.3km | 48°44.635’ N | 3°56.722’ W |
| *Laminaria digitata* | France | Roscoff | Stn4 - incoming | 4km | 48°45.023’ N | 3°56.593’ W |
| *Laminaria digitata* | France | Roscoff | Stn1 - outgoing | 0km | 48°42.896’ N | 3°57.353’ W |
| *Laminaria digitata* | France | Roscoff | Stn2 - outgoing | 1.5km | 48°43.641’ N | 3°57.079’ W |
| *Laminaria digitata* | France | Roscoff | Stn3 - outgoing | 3.3km | 48°44.661’ N | 3°56.716’ W |
| *Laminaria digitata* | France | Roscoff | Stn4 - outgoing | 4km | 48°45.047’ N | 3°56.641’ W |
| *Lessonia trabeculata* | Chile | Las Cruces | Stn1 | 0.7km | 33°29'24.64" S | 71°39'4.47" W |
| *Lessonia trabeculata* | Chile | Las Cruces | Stn2 | 2km | 33°29'21.45" S | 71°39'53.85" W |
| *Lessonia trabeculata* | Chile | Las Cruces | Stn3 | 3km | 33°29'26.20" S | 71°40'39.35" W |
| *Lessonia trabeculata* | Chile | Las Cruces | Stn4 | 4.8km | 33°29'6.41" S | 71°41'41.07" W |
| *Lessonia trabeculata* | Chile | Algarrobo | Stn1 | 0.5km | 33°21'22.13" S | 71°41'34.32" W |
| *Lessonia trabeculata* | Chile | Algarrobo | Stn2 | 1.5km | 33°20'56.05" S | 71°42'4.82" W |
| *Lessonia trabeculata* | Chile | Algarrobo | Stn3 | 2km | 33°21'31.73" S | 71°42'38.74" W |
| *Lessonia trabeculata* | Chile | Algarrobo | Stn4 | 3.3km | 33°22'5.80" S | 71°43'25.40" W |
| *Saccharina latissima* | Northern Ireland | Rathlin | upstream | 0.1km | 55°17.225’N | 6°13.555’ W |

| | | | | | | |
|---|---|---|---|---|---|---|
| *Saccharina latissima* | Northern Ireland | Rathlin | upstream | 0.1km | 55°17.345’N | 6°13.551’ W |
| *Saccharina latissima* | Northern Ireland | Rathlin | kelp farm | 0km | 55°17.364’N | 6°13.481’ W |
| *Saccharina latissima* | Northern Ireland | Rathlin | kelp farm | 0km | 55°17.368’ N | 6°13.460’ W |
| *Saccharina latissima* | Northern Ireland | Rathlin | kelp farm | 0km | 55°17.371’N | 6°13.412’ W |
| *Saccharina latissima* | Northern Ireland | Rathlin | downstream | 0.2km | 55°17.375’ N | 6°13.346’ W |
| *Saccharina latissima* | Northern Ireland | Rathlin | downstream | 0.2km | 55°17.389’ N | 6°13.305’ W |
| *Saccharina latissima* | Northern Ireland | Rathlin | downstream | 0.2km | 55°17.396’ N | 6°13.281’ W |
| *Saccharina latissima* | Northern Ireland | Rathlin | downstream | 0.2km | 55°17.406’ N | 6°13.256’ W |
| *Fucus vesiculosus* | Finland | Tvärminne | Stn1 | 0km | 59°48.849' N | 23°14.802' E |
| *Fucus vesiculosus* | Finland | Tvärminne | Stn2 | 1km | 59°48.643' N | 23°13.835' E |
| *Fucus vesiculosus* | Finland | Tvärminne | Stn3 | 2.3km | 59°48.327' N | 23°12.529' E |
| *Fucus vesiculosus* | Finland | Tvärminne | Stn4 | 3.4km | 59°48.107' N | 23°11.502' E |
| *Fucus vesiculosus* | Finland | Tvärminne | Stn5 | 5km | 59°47.745' N | 23°9.907' E |
| *Sargassum fluitans* | United States | Western North Atlantic | Stn21 | 0km | 33.740° N | 75.291° W |
| *Sargassum fluitans* | United States | Western North Atlantic | Stn22 | 2100km | 42.86173° N | 53.86997° W |
| *Sargassum fluitans* | United States | Western North Atlantic | Stn23 | 500km | 34°04'59.0"N | 70°05'52.0"W |
| *Sargassum fluitans* and *natans* | Mexico | Puerto Morelos | Patch | inside and outside | 20°49'56.89"N | 86°52'15.41"W |

1015

**Table S2.** Overview of species, sampling time, location, including latitude and longitude and incubation volume of all six different mesocosm experiments.

| Species | Sampling time | Location | Latitude | Longitude | Incubation volume |
|---|---|---|---|---|---|
| *Ecklonia maxima* | March 2022 | South Africa, Southern Atlantic | 33°55.8' S | 18°22.8' E | 220 L |
| *Ecklonia radiata* | May 2022 | New Zealand, Southern Pacific | 37°40.3' S | 176°10.0' E | 220 L |
| *Laminaria digitata* | August 2022 | France, The English Channel | 48°43.6' N | 71°38.0' W | 250 L |
| *Lessonia trabeculata* | January 2022 | Chile, Southern Pacific | 33°30.1' S | 71°38.0' W | 250 L |
| *Fucus vesiculosus* | October 2022 | Finland, Baltic Sea | 59°50.7' N | 23°14.9' E | 57 L |
| *Sargassum fluitans* | May 2022 | United States of America, Sargasso Sea | 35°11.8' N | 70°0.3' W | 40 L |
